# ASCT2 is the primary serine transporter in cancer cells

**DOI:** 10.1101/2023.10.09.561530

**Authors:** Kelly O. Conger, Christopher Chidley, Mete Emir Ozgurses, Huiping Zhao, Yumi Kim, Svetlana E. Semina, Philippa Burns, Vipin Rawat, Ryan Sheldon, Issam Ben-Sahra, Jonna Frasor, Peter K. Sorger, Gina M. DeNicola, Jonathan L. Coloff

## Abstract

The non-essential amino acid serine is a critical nutrient for cancer cells due to its diverse biosynthetic functions. While some tumors can synthesize serine *de novo*, others are auxotrophic for serine and therefore reliant on the uptake of exogenous serine. Importantly, however, the transporter(s) that mediate serine uptake in cancer cells are not known. Here, we characterize the amino acid transporter ASCT2 (coded for by the gene *SLC1A5*) as the primary serine transporter in cancer cells. ASCT2 is well-known as a glutamine transporter in cancer, and our work demonstrates that serine and glutamine compete for uptake through ASCT2. We further show that ASCT2-mediated serine uptake is essential for purine nucleotide biosynthesis and that ERα promotes serine uptake by directly activating *SLC1A5* transcription. Together, our work defines an additional important role for ASCT2 as a serine transporter in cancer and evaluates ASCT2 as a potential therapeutic target in serine metabolism.

## INTRODUCTION

Amino acid metabolism is central to a cell’s ability to accumulate biomass and proliferate^1,2^. In particular, the non-essential amino acid L-serine (serine) has received considerable attention, in part because of its importance as a precursor for the synthesis of numerous macromolecules including proteins, lipids, and nucleotides^3–8^. As a non-essential amino acid, serine can be synthesized via the serine synthesis pathway or consumed through dietary protein^6^. It has been observed that some cancer cells increase serine synthesis pathway flux by increasing expression of the synthesis pathway genes *PHGDH, PSAT1,* and *PSPH*^9,10^. These and other observations motivated the development of PHGDH inhibitors as potential cancer therapies^11^. However, in most cases uptake of exogenous serine appears to be sufficient to offset inhibition of the synthesis pathway^12,13^, thus limiting the efficacy of PHGDH inhibitors as potential therapeutic agents.

The importance of exogenous serine uptake has prompted investigations into serine starvation as an alternative approach of targeting serine metabolism in cancer^14^. Deprivation of serine can be achieved *in vivo* through dietary serine and glycine restriction, which reduces circulating serine levels by ∼50%^15–17^. Previous work has identified several genetic alternations that increase sensitivity to dietary serine starvation^15–17^, as well as how dietary serine deprivation can sensitize cancer cells to other therapies^16,18,19^. Importantly, cancer cells that are auxotrophic for serine are particularly sensitive to exogenous serine starvation both *in vitro* and *in vivo*^17,20^. Tumors that are naturally auxotrophic for serine, including luminal/estrogen receptor positive (ER+) breast tumors^20^, are prime candidates for serine deprivation therapy^17,21–23^. While dietary serine starvation is a promising therapeutic approach, it is possible that long-term systemic serine deficiency has the potential to cause peripheral neuropathy^24–29^. It has also recently been observed that stromal cells in the tumor microenvironment can secrete serine for uptake by auxotrophic cancer cells, which could also limit the efficacy of serine starvation therapy^23^. Another potential approach of starving cancer cells of serine that could be used alone or in combination with serine starvation is preventing serine entry into cancer cells.

However, we still do not know which transporter(s) contribute to serine uptake in cancer. Initial *in* vitro characterization of substrate specificity for amino acid transporters using liposomes or *Xenopus leavis* oocytes has identified six transporters - ASCT1, ASCT2, SNAT1, SNAT2, LAT1, and LAT2 – as being capable of transporting serine^30–34^. However, apart from the identification of ASCT1 as a mediator of serine uptake in the brain^35–38^, these transporters have not been investigated for potential contributions to serine uptake in any physiological conditions, including in cancer cells. Recently, the novel serine transporter SFXN1 has been characterized for its contributions to mitochondrial serine transport^39^, but it is unlikely to contribute to extracellular serine uptake due to its localization at the mitochondria.

Here, we present evidence that the amino acid transporter ASCT2 (coded by the gene *SLC1A5*) is the primary serine transporter in luminal/ER+ breast cancer and other tumor cell lines. While ASCT2 has been extensively studied as a glutamine transporter in cancer, our work is the first to describe a fundamental role for ASCT2 in cancer cell serine metabolism. Although ASCT2 contributes to serine uptake in all cancer cell lines tested, we find that ASCT2-mediated serine uptake is particularly important in serine auxotrophic cells and in limited serine conditions. Additionally, we demonstrate that serine provided by ASCT2 is critical for purine nucleotide biosynthesis and discover a novel mechanism of *SLC1A5* regulation by ERα in ER+ breast cancer. Finally, we establish that ASCT2 may be a viable therapeutic target in luminal breast cancer, particularly in combination with dietary serine starvation. This work expands our understanding of serine metabolism and amino acid transporter biology in cancer and has important implications in the effort to target serine metabolism for cancer therapy.

## RESULTS

### Identification of ASCT2 as a serine transporter in luminal/ER+ breast cancer cells

We have recently discovered that luminal/ER+ breast cancer cells are auxotrophic for serine^20^ and reasoned that their dependence on exogenous serine would make them an ideal model for identifying serine transporters. Because there are numerous transporters that have the potential to transport serine, we made use of a two-part CRISPRi and CRISPRa screen with a library targeting 489 human transporter genes belonging to the SLC and ABC families^40^. This library has recently been used to identify transporters for a variety of amino acids, but these efforts were less successful for identifying serine transporters given that they were screened in cells that are not auxotrophic for serine^40^. Using this library, we therefore utilized our pooled screening approach to construct single transporter knockdown or overexpression cells in the serine-auxotrophic luminal/ER+ breast cancer cell line MCF7. To focus our screen on serine transporters and not just essential transporter genes, we compared the impact of SLC and ABC gene perturbation in (i) low serine conditions (50 μM for CRISPRi and 18 μM for CRISPRa), which inhibit growth by approximately 25-75% (data not shown), and in (ii) serine conditions (285 μM) found in the commonly used tissue culture medium RPMI-1640 (RPMI). This strategy allows the identification of transporters that contribute to serine uptake in MCF7 cells at endogenous expression levels (CRISPRi) and also of transporters that have the ability to transport serine when over-expressed (CRISPRa) (Figure 1A).

**Figure 1.**
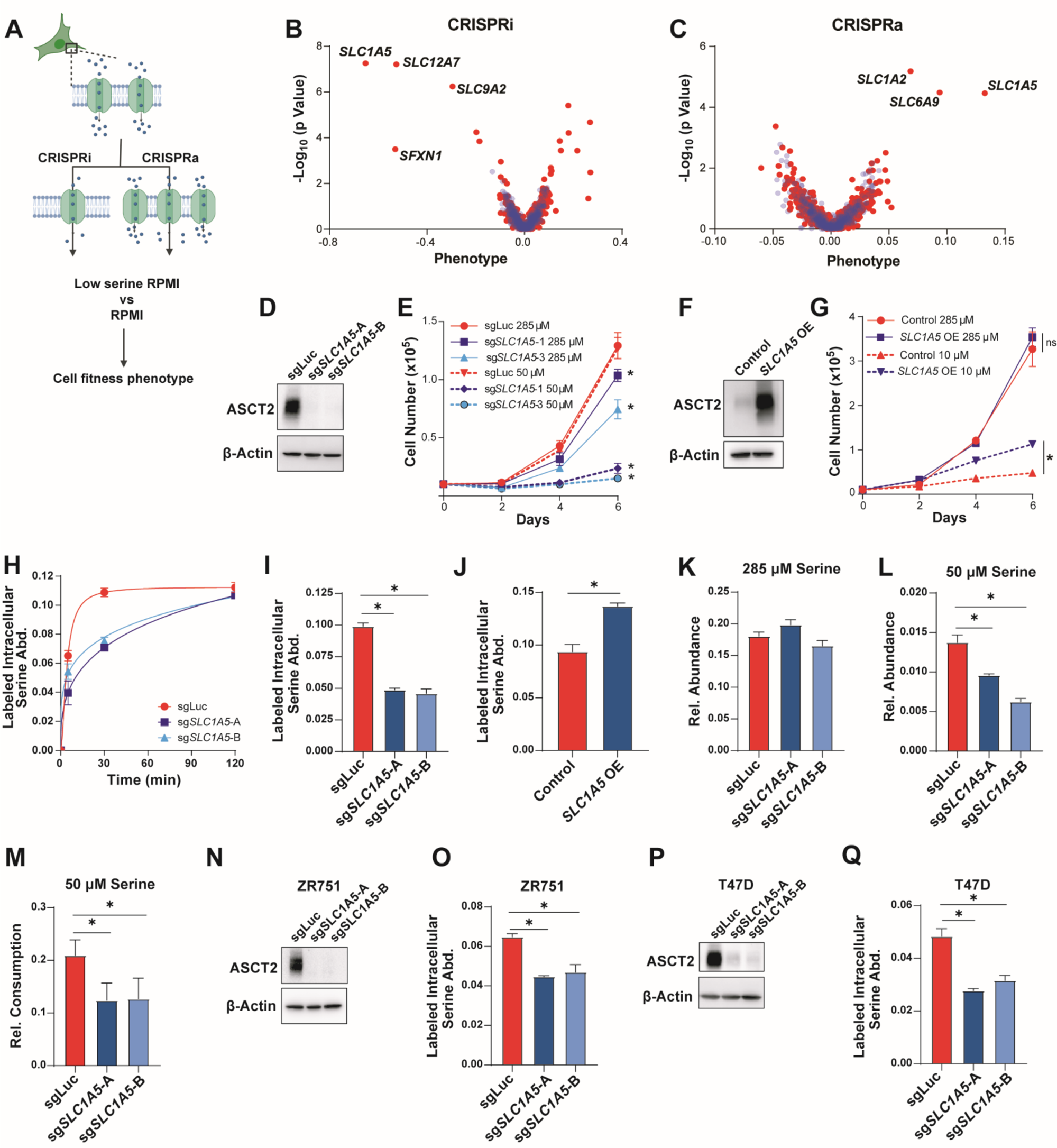
Identification of ASCT2 as a serine transporter in luminal breast cancer cells. A). Schematic of the CRISPRi/CRISPRa screening method. B & C). Results of the CRISPRi (B) and CRISPRa (C) screen performed in RPMI with low serine. Blue dots are pseudogenes that represent the technical noise of the assay, red dots represent transporter genes. D). Representative western blot of control (sgLuc) and ASCT2 KO (sg*SLC1A5*-A and sg*SLC1A5*-B) MCF7 cells. E). Growth assay of MCF7 control (sgLuc) or ASCT2 KO (sg*SLC1A5*) cells in complete (285 µM serine) RPMI (solid lines) or low serine (50 µM serine) RPMI (dotted lines). Values are the means ±SD of triplicate samples from an experiment representative of 3 independent experiments. *p < 0.05 by two-way ANOVA. MCF7 sgLuc is compared to sg*SLC1A5* for each serine dose. F). Representative western blot of control and ASCT2 over-expressing (*SLC1A5* OE) MCF7 cells. G). Growth assay of control and ASCT2 over-expressing (*SLC1A5* OE) MCF7 cells in complete (285 µM serine) RPMI (solid lines) or low serine (10 µM serine) RPMI (dotted lines). Values are the mean ±SD of triplicate samples. *p < 0.05 by two-way ANOVA. H). Accumulation of U-^13^C_3_-^15^N-serine in MCF7 sgLuc and sg*SLC1A5* cells over time. Values are the means ±SD of triplicate samples from an experiment representative of 2 independent experiments. I). Acute serine uptake in MCF7 control (sgLuc) and ASCT2 KO (sg*SLC1A5*) cells in complete media. Values are the mean ±SD of triplicate samples from an experiment representative of 3 independent experiments. *p < 0.05 by Welch’s t-test. J). Acute serine uptake in MCF7 control and ASCT2 over-expressing (*SLC1A5* OE) cells in complete media. Values are the mean ±SD of triplicate samples from an experiment representative of 3 independent experiments. *p < 0.05 by Welch’s t-test. K). Intracellular serine abundance in MCF7 control (sgLuc) and ASCT2 KO (sg*SLC1A5*) cells cultured in complete media. Values are the mean ±SD of triplicate samples from an experiment representative of 2 independent experiments. L). Intracellular serine abundance in MCF7 control (sgLuc) and ASCT2 KO (sg*SLC1A5*) cells after exposure to low serine (50 µM) conditions for six hours. Values are the mean ±SD of triplicate samples from an experiment representative of 2 independent experiments. *p < 0.05 by Welch’s t-test. M). Net consumption of serine from the medium by MCF7 control (sgLuc) and ASCT2 KO (sg*SLC1A5*) cells after exposure to low serine (50 µM) conditions for six hours. Values are the mean ±SD of triplicate samples from an experiment representative of 2 independent experiments. *p < 0.05 by Welch’s t-test. N). Representative western blot of control (sgLuc) and ASCT2 KO (sg*SLC1A5*) ZR751 cells. O). Acute serine uptake in ZR751 control and ASCT2 KO (sg*SLC1A5*) cells in complete media. Values are the mean ±SD of triplicate samples from an experiment representative of 3 independent experiments. *p < 0.05 by Welch’s t-test. P). Representative western blot of control (sgLuc) and ASCT2 KO (sg*SLC1A5*) T47D cells. Q). Acute serine uptake in T47D control and KO (sg*SLC1A5*) cells in complete media. Values are the mean ±SD of triplicate samples from an experiment representative of 3 independent experiments. *p < 0.05 by Welch’s t-test.

Using this approach, we identified *SLC1A5*, which codes for the amino acid transporter ASCT2, as the top scoring hit in both the CRISPRi and CRISPRa screens (Figures 1B, C). Another significant hit in our CRISPRi screen was *SFXN1*, which was recently shown to be a mitochondrial serine transporter^39^, demonstrating that our screening approach is capable of identifying transporters involved in serine metabolism. *SLC12A7*^41^ and *SLC9A2*^42^ also scored in our CRISPRi screen (Figure 1B), but their specificity as ion transporters suggests that their phenotypes likely arise via indirect effects on serine metabolism. *SLC1A2* (glutamate)^43^ and *SLC6A9* (glycine)^44^ were also hits in our CRISPRa screen (Figure 1C) but it is unclear whether these transporters directly contribute to serine uptake or rather provide other amino acids that support growth in low-serine conditions.

To validate ASCT2’s role as a serine transporter, we constructed *SLC1A5* MCF7 knockout (KO) cells using CRISPR/Cas9 and two unique sgRNAs (Figure 1D) where we found that loss of ASCT2 only slightly reduces cell growth in normal RPMI (high levels of serine) but strongly reduces proliferation at lower levels of extracellular serine (50 μM) (Figures 1E, S1A, S1B). Conversely, over-expression of ASCT2 significantly rescues growth in very low serine (10 μM) (Figures 1F-G, S1C). To directly test whether loss of ASCT2 affects serine uptake, we treated control and KO cell lines with media containing U-^13^C_3_-^15^N-serine and measured the accumulation of intracellular labeled serine over time by GC-MS, which revealed a significant defect in serine uptake in ASCT2 KO cells (Figure 1H). Because intracellular labeled serine plateaued by 30 minutes in control cells (Figure 1H), for all future serine uptake assays we measured intracellular labeled serine abundance after four minutes. This approach revealed that ASCT2 KO cells exhibit decreased acute serine uptake (Figure 1I), while over expression of ASCT2 significantly increases serine uptake in MCF7 cells (Figure 1J). Notably, we did not observe differences in total serine abundance in ASCT2 KO cells in full serine conditions (Figure 1K), however, ASCT2 KO cells display decreased serine abundance after six hours of culture at low (50 μM) serine (Figure 1L). ASCT2 KO cells also consume less serine (as measured by serine depletion from the medium) when cultured in 50 μM serine (Figure 1M). Knockout of ASCT2 in two other luminal breast cancer cell lines revealed a similar decrease in acute serine uptake (Figures 1N-Q). Together, these results demonstrate that ASCT2 is a major contributor to serine uptake in luminal breast cancer cells.

### ASCT2 contributes to serine uptake but does not sensitize to low serine in non-auxotrophic cells

While serine auxotrophic cell lines require uptake of exogenous serine, non-auxotrophic cells can synthesize serine *de novo* and are less sensitive to low serine conditions. To determine whether ASCT2 contributes to serine uptake in non-auxotrophic cell lines, we knocked out ASCT2 in the basal/triple-negative breast cancer cell lines HCC1806 and SUM149 that we have previously shown to be non-auxotrophic for serine^20^ (Figures 2A, S2A). Similar to the luminal/ER+ lines, we found that HCC1806 and SUM149 cells lacking ASCT2 display decreased serine uptake (Figures 2B-C). We also knocked out ASCT2 in A498 clear cell renal cell carcinoma (ccRCC) cells and A549 and Calu6 lung cancer cell lines and saw decreased serine uptake in all cases (Figures 2D-F, S2B-D), demonstrating that ASCT2 contributes to serine uptake even in non-auxotrophic cell lines and cell lines from diverse tumor types. While loss of ASCT2 does reduce proliferation of HCC1806 cells in full serine conditions (likely due to reduced glutamine uptake^45^), it does not sensitize to low serine (Figure 2G). Importantly, however, preventing *de novo* serine biosynthesis by knocking out *PSAT1*^20^ does sensitize HCC1806 cells to loss of ASCT2 in low serine conditions (Figures 2H-I). Additionally, loss of ASCT2 in HCC1806 cells also leads to increased serine biosynthesis from glucose (Figure 2J), further indicating the importance of ASCT2 in serine metabolism. Finally, overexpression of PSAT1 in MCF7 cells partially rescues growth in low serine after loss of ASCT2 (Figures 2K-L). Together, these data suggest that ASCT2 acts as a serine transporter in both auxotrophic and non-auxotrophic cell lines, but only auxotrophic cells rely on ASCT2 to support growth in low serine conditions.

**Figure 2.**
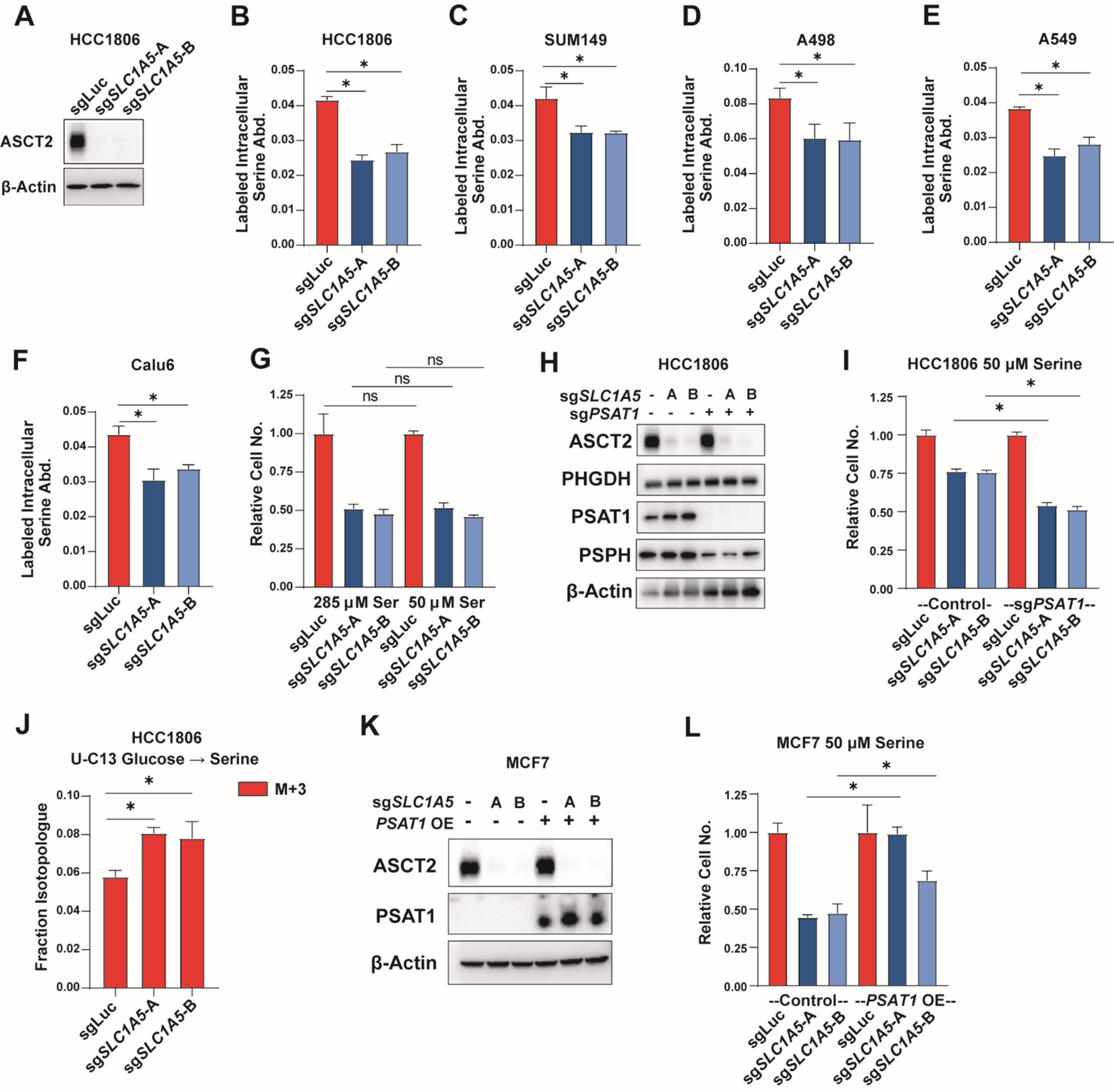
ASCT2 contributes to serine uptake in non-auxotrophic cells. A). Representative western blot of control (sgLuc) and ASCT2 KO (sg*SLC1A5*-A and sg*SLC1A5*-B) HCC1806 cells. B). Acute serine uptake in HCC1806 control (sgLuc) and ASCT2 KO (sg*SLC1A5*) in complete media. Values are the mean ±SD of triplicate samples from an experiment representative of 3 independent experiments. *p < 0.05 by Welch’s t-test. C-F). Acute serine uptake in SUM149 (D), A498 (E), A549 (F), and Calu6 (G) control (sgLuc) and ASCT2 KO (sg*SLC1A5*) cell lines at complete RPMI concentration (285 µM). Values are the mean ±SD of triplicate samples from an experiment representative of 3 independent experiments. *p < 0.05 by Welch’s t-test. G). Growth assay of HCC1806 control (sgLuc) or ASCT2 KO (sg*SLC1A5*) cells grown in complete RPMI (285 µM) or low serine (50 µM) RPMI. Values are the mean ±SD of triplicate samples from an experiment representative of 3 independent experiments *p < 0.05 by Welch’s t-test. H) Representative western blot of HCC1806 with either ASCT2 KO alone (sg*SLC1A5*), or in combination with KO of PSAT1 (sg*PSAT1*). I) Growth assay of HCC1806 ASCT2 KO alone and ASCT2/PSAT1 double KO cells grown in low serine (50 µM) RPMI. Values are the mean ±SD of triplicate samples from an experiment representative of 3 independent experiments. *p < 0.05 by Welch’s t-test. J). *De novo* serine biosynthesis from glucose in HCC1806 control (sgLuc) and ASCT2 KO (sg*SLC1A5*) cells for 24 hours. Values are the means ±SD of M+3 labeled serine from triplicate samples from an experiment representative of 2 independent experiments. *p < 0.05 by Welch’s t-test. K). Representative western blot of MCF7 cells with either ASCT2 KO alone or in combination with PSAT1 overexpression. L). Growth assay in low serine (50 µM) RPMI of MCF7 ASCT2 single KO, PSAT1 OE, and combined ASCT2 KO/PSAT1 OE. Values are the mean ±SD of triplicate samples from an experiment representative of 2 independent experiments. *p < 0.05 by Welch’s t-test.

### Transporter compensation in the absence of ASCT2

**L**oss of ASCT2 reduces but does not eliminate serine uptake suggesting that there are secondary transporters that also contribute to serine uptake. We interrogated potential secondary transporters based on those published as capable of transporting serine in *Xenopus* oocytes or liposomes: ASCT1 (*SLC1A4*)^35–38^, SNAT1 (*SLC38A1*)^34^, SNAT2 (*SLC38A2*)^46^, LAT1 (*SLC7A5*)^32^, LAT2 (*SLC7A8*)^31^ and ATB0+ (*SLC6A14*)^47^. First, we evaluated the gene expression of these candidate transporters upon serine and glycine starvation, which we reasoned might be induced if they are involved in serine uptake. With the exception of *SLC7A8*, all of the candidate transporters are upregulated upon serine and glycine deprivation, with *SLC1A4* (which encodes the related transporter ASCT1) showing the most significant increase (Figure 3A). We also evaluated the protein expression of these transporters in ASCT2 KO cells to determine if their expression levels might change to compensate for loss of ASCT2. Here, we found that only ASCT1 is upregulated in ASCT2 KO MCF7 cells (Figures 3B). To evaluate a role for ASCT1 in serine uptake, we generated ASCT1 and ASCT2 single and double KO MCF7 cells (Figure 3C). Importantly, while loss of ASCT1 alone slightly reduces serine uptake, loss of both transporters does not significantly reduce uptake beyond what is observed with ASCT2 alone (Figure 3E). Further, overexpression of ASCT1 to a similar degree as seen in ASCT2 KO cells induces a slight, but significant, increase in serine uptake (Figures 3F, 3G). However, combined loss of ASCT1 and ASCT2 does not dramatically reduce proliferation in low serine (Figure 3D), nor does ASCT1 overexpression rescue proliferation in low serine in ASCT2 KO cells (Figure 3H). These results suggest that ASCT1 may contribute to serine uptake in MCF7 cells, but its contribution is small relative to that of ASCT2.

**Figure 3.**
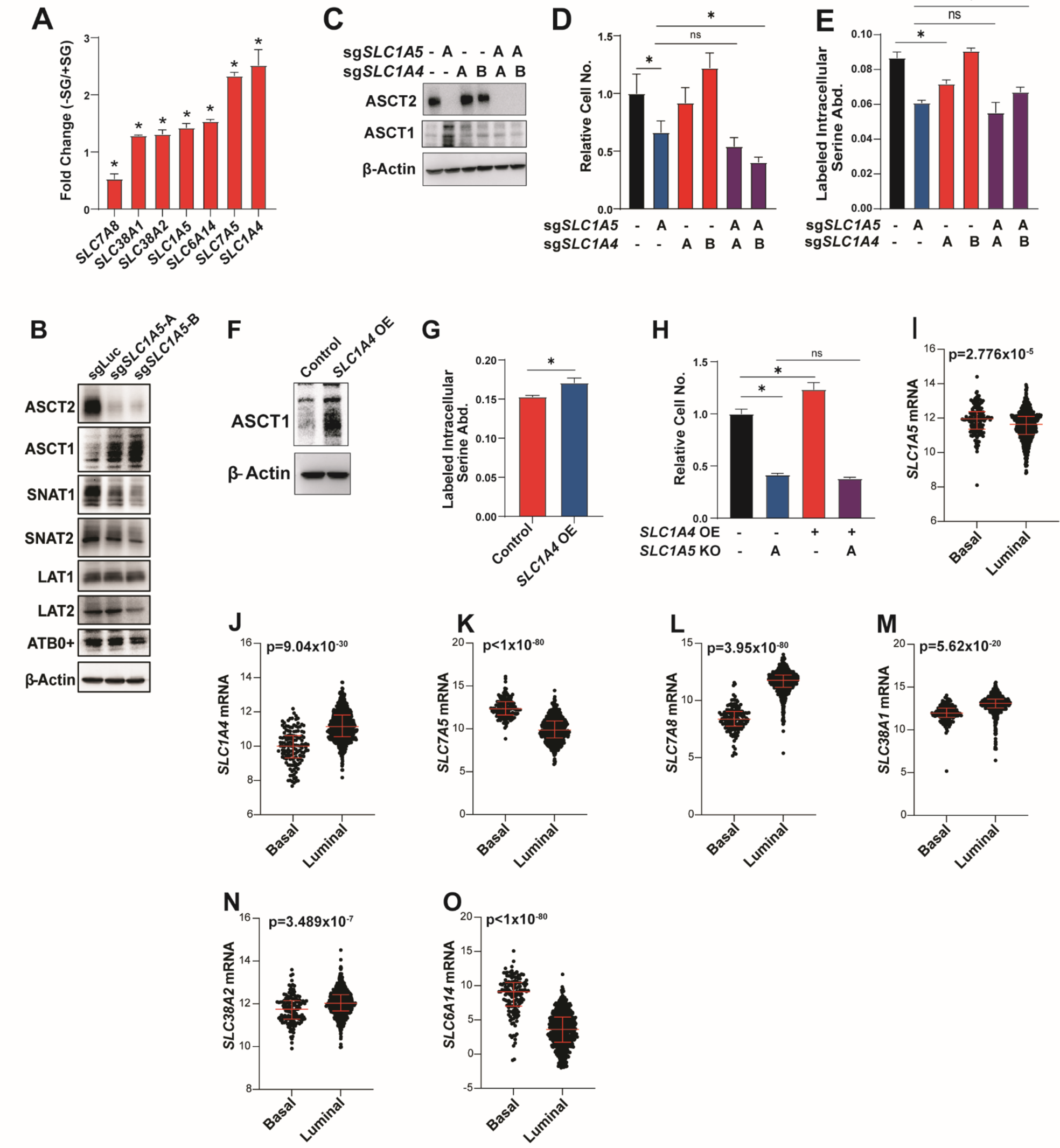
Transporter compensation in the absence of ASCT2. A). RNASeq from MCF7 cells grown in the presence or absence of serine and glycine for 48 hours prior to harvesting RNA. Values represent the fold change difference in -SG vs +SG conditions for triplicate samples. B). Western blot from MCF7 control (sgLuc) and ASCT2 KO (sg*SLC1A5*) cells. C). Representative western blot from single and double ASCT1 (sg*SLC1A4*) and ASCT2 (sg*SLC1A5*) KO MCF7 cells. D). Growth assay in low (50 µM) serine for single and double ASCT1 (sg*SLC1A4*) and ASCT2 (sg*SLC1A5*) KO MCF7 cells. Values are the mean ±SD of triplicate samples from an experiment representative of 2 independent experiments. *p < 0.05 by Welch’s t-test. E). Acute serine uptake assay from single and double ASCT1 (sg*SLC1A4*) and ASCT2 (sg*SLC1A5*) KO MCF7 cells. Values are the mean ±SD of triplicate samples from an experiment representative of 2 independent experiments. *p < 0.05 by Welch’s t-test. F). Representative western blot for control and ASCT1 over-expressing (*SLC1A4* OE) MCF7 cells. G). Acute serine uptake from control and ASCT1 over-expressing (*SLC1A4* OE) MCF7 cells in complete RPMI media. Values are the mean ±SD of triplicate samples from an experiment representative of 2 independent experiments. *p < 0.05 by Welch’s t-test. H). Growth assay in ASCT2 KO (sg*SLC1A5*) MCF7 cells with or without ASCT1 over-expression (*SLC1A4* OE) grown at low (50 µM) serine. Values are the mean ±SD of triplicate samples from an experiment representative of 3 independent experiments. *p < 0.05 by Welch’s t-test. I-O). Transporter gene expression by breast tumor subtype in the TCGA pan-cancer atlas dataset.

To evaluate contributions to serine uptake from the other five candidate transporters we generated single and double transporter KOs with ASCT2 (Figures S3A, S3C, SE, S3G, S3I), where only loss of LAT1 on its own showed a slight reduction in serine uptake, and none of the double knockout cells exceeded the effects seen with ASCT2 alone (Figures S3B, S3D, S3F, S3H, S3J). Given these results, we feel that our data is most consistent with a model in which ASCT2 is the primary serine transporter in cancer cells, but a network of secondary transporters can maintain serine uptake in the absence of ASCT2, particularly in high serine conditions. Consistent with this model, all six potential serine transporter genes are expressed in luminal breast tumors, with *SLC1A4*, *SLC38A1*, *SLC38A2*, and *SLC7A8* being enriched in luminal breast tumors relative to basal (Figures 3I-O).

### Serine and glutamine compete for uptake by ASCT2

While ASCT2 has not previously been described as a serine transporter in cancer, it has been extensively characterized as a glutamine transporter^45,48–51^. Indeed, we find that knockout and overexpression of ASCT2 in MCF7 cells causes a decrease and increase, respectively, in acute glutamine uptake (Figure 4A-B), and that loss of ASCT2 sensitizes MCF7 cells to decreasing glutamine concentrations (Figures 4C-D). To further examine ASCT2 substrate dynamics, we starved MCF7 cells of glutamine for one hour and found that intracellular serine abundance is strongly increased upon glutamine starvation (Figure 4E). Further, we find that the accumulation of intracellular serine after glutamine starvation is delayed in ASCT2 KO cells (Figure 4F). These data are suggestive of a model where glutamine and serine compete for uptake through ASCT2. To test this model, we performed serine and glutamine uptake assays in cells treated with lower or higher concentrations of serine and glutamine. Not surprisingly, we found that by raising the serine concentration to match RPMI glutamine levels (2.0 mM Ser), serine uptake is increased (Figure 4G). Interestingly, however, lowering glutamine levels by a factor of 10 (0.2 mM Gln) was also sufficient to increase serine uptake (Figure 4G), and inverting the ratio of serine to glutamine (2.0 mM Ser, 0.2 mM Gln), incudes the greatest increase in serine uptake (Figure 4G).

**Figure 4.**
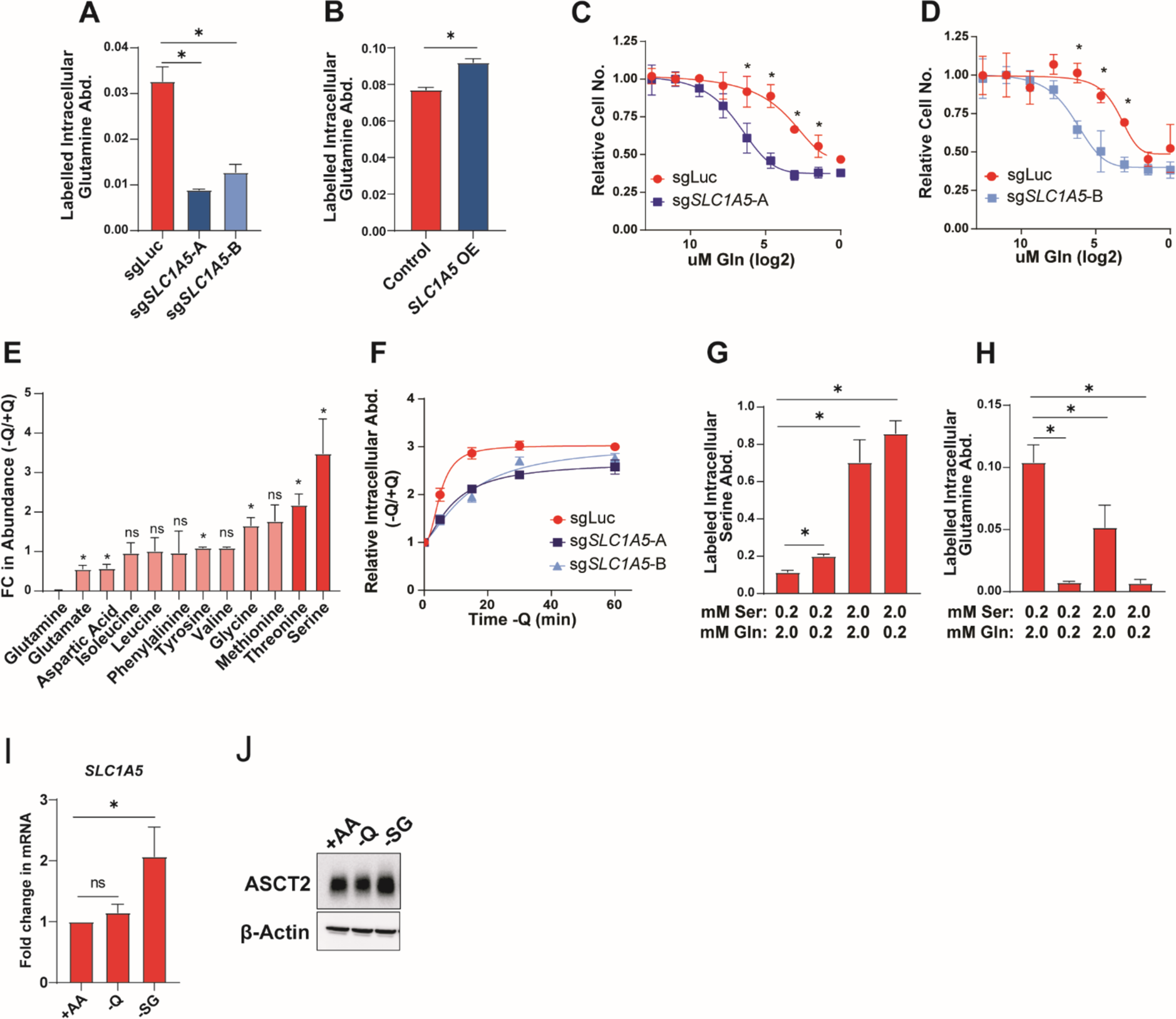
Serine and glutamine compete for uptake by ASCT2. A). Acute glutamine uptake in MCF7 control (sgLuc) or *SLC1A5* KO (sg*SLC1A5*) cells. Values are the mean ±SD of triplicate samples from an experiment representative of 3 independent experiments. *p < 0.05 by Welch’s t-test. B). Acute glutamine uptake in MCF7 control or *SLC1A5* over-expressing (*SLC1A5* OE) cells. Values are the mean ±SD of triplicate samples from an experiment representative of 2 independent experiments. *p < 0.05 by Welch’s t-test C-D). Glutamine dose curve with MCF7 control (sgLuc) and sg*SLC1A5* cells. Values are the means ±SD of triplicate samples from an experiment representative of 3 independent experiments. *p < 0.05 by Welch’s t-test. Relative cell counts are normalized to each group’s count at normal RPMI glutamine dose. E). Intracellular amino acid abundance from MCF7 cells cultured in the absence of glutamine for one hour. Values are the mean ±SD of the relative abundance for each amino acid in -Q relative to +Q conditions from an experiment representative of 2 independent experiments. *p < 0.05 by Welch’s t-test comparing -Q to +Q conditions. F). Intracellular serine abundance from MCF7 control (sgLuc) and *SLC1A5* KO (sg*SLC1A5*) cells cultured in the absence of glutamine for 5, 15, 30, and 60 minutes. Values are the mean ±SD of the relative serine abundance for each group at each time point. G-H). Acute serine (G) and glutamine (H) uptake in MCF7 cells cultured in the indicated concentrations of serine and glutamine. Values are the mean ±SD of triplicate samples from an experiment representative of 3 independent experiments. *p < 0.05 by Welch’s t-test. I). qPCR data for *SLC1A5* expression in MCF7 cells cultured in complete RPMI (+AA), glutamine deprivation (-Q), or serine and glycine deprivation (-SG) for 48 hours. Values are the means ±SD of the fold change in *SLC1A5* mRNA from three independent experiments. *p < 0.05 by Welch’s t-test. J). Representative western blot of MCF7 cells cultured in complete RPMI (+AA), glutamine deprivation (-Q), or serine and glycine deprivation (-SG) for 48hours.

Conversely, we found that glutamine uptake is significantly reduced when either glutamine is lowered or serine is increased, or both (Figure 4H). Together, we interpret these data as in support of a model in which serine and glutamine are competitive substrates of ASCT2. Interestingly, *SLC1A5* and ASCT2 expression are induced in response to deprivation of serine, but not glutamine (Figures 4I, 4J), further supporting a prominent role for ASCT2 in serine metabolism.

### ASCT2 is required for purine nucleotide biosynthesis when serine levels are limited

Serine is important for several downstream pathways (Figure 5A) and has received considerable attention for its role as a precursor for purine nucleotide and antioxidant biosynthesis^4,52,53^. These contributions primarily occur through the action of serine hydroxymethyltransferase (SHMT1/2), which converts serine to glycine and one carbon units that can be used in glutathione (GSH) and purine nucleotide synthesis (Figure 5B). By tracing the incorporation of serine-derived carbon (U-^13^C_3_-serine) into downstream metabolites, we observed that loss of ASCT2 results in decreased generation of glycine from serine in both complete and low serine conditions (Figure 5C). And while loss of ASCT2 does not strongly affect the synthesis of GSH or the purine nucleotides inosine monophosphate (IMP), adenosine monophosphate (AMP), and guanosine monophosphate (GMP) in high serine, GSH and purine nucleotide biosynthesis are significantly reduced in ASCT2 KO cells in low serine conditions (Figures 5D – 5G). Importantly, because ASCT2 KO cells proliferate more slowly in low serine, the decreased biosynthesis observed in these conditions could be either a cause or a consequence of reduced proliferation. To address this, we provided cells with hypoxanthine, a substrate for the purine salvage pathway and/or the antioxidant N-acetyl cysteine (NAC), both of which have been used previously to rescue the effects of serine starvation (Figure 5A)^19,54^. Addition of hypoxanthine was sufficient to strongly (but not completely) rescue the growth of ASCT2 KO cells cultured in low serine, while NAC had no effect either alone or in combination with hypoxanthine (Figure 5H). These results indicate that loss of ASCT2 in low serine impairs purine biosynthesis, which results in decreased proliferation.

**Figure 5.**
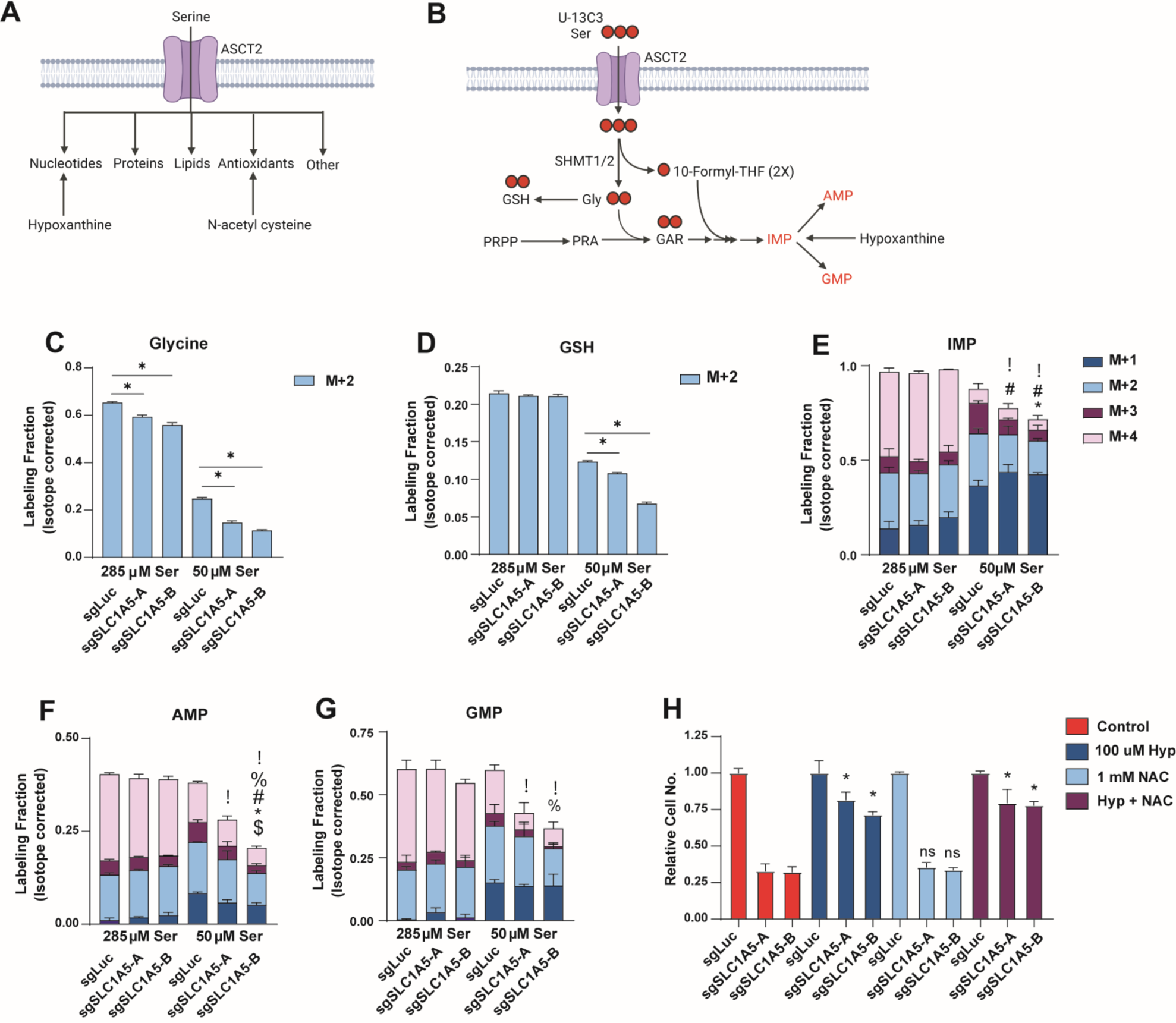
ASCT2 is required for purine nucleotide biosynthesis when serine levels are limited. A). Schematic illustrating how serine is utilized inside the cell, and how nucleotide and antioxidant metabolic pathways can be rescued in the absence of serine. B). Schematic illustrating how U-^13^C_3_ -serine can be traced through the purine and GSH biosynthesis pathways. C-G). Stable isotope tracing results for MCF7 control (sgLuc) and *SLC1A5* KO (sg*SLC1A5*) cells cultured in media containing U-^13^C_3_-serine at normal or low serine concentrations for eight hours. Values are the mean ±SD of triplicate samples. P<0.05 by Welch’s t-test for each isotopomer in sg*SLC1A5* compared to sgLuc (! = M+0, % = M+1, # = M+2, * = M+3, $ = M+4). H). Growth assay of MCF7 control (sgLuc) and *SLC1A5* KO (sg*SLC1A5*) cells cultured as indicated. Values are the means ±SD of triplicate samples representative of three independent experiments. *p < 0.05 by Welch’s t-test.

### ERα promotes serine uptake via direct regulation of *SLC1A5*

Most luminal breast tumors express estrogen receptor α (ERα) and are dependent on it for growth and proliferation^55^. Because ERα is known to control many essential growth pathways, including several metabolic pathways and amino acid transporters^56–58^, we hypothesized that estrogen signaling might also promote ASCT2 expression and serine uptake. Indeed, we found that treatment of MCF7 cells with the ERα inhibitors tamoxifen and fulvestrant (ICI) lowers ASCT2 (Figure 6A), and *SLC1A5* (Figure 6B) expression, and inhibits serine uptake (Figures 6C). Using the EstrogeneDB database (www.estrogene.org), which compiles microarray and RNAseq studies of estrogen-regulated gene expression, we observed that estrogen regulates *SLC1A5* mRNA expression in multiple ER+ cell lines at various doses and treatment durations (Figure S4A). Additionally, we found that estrogen starvation significantly reduces ASCT2 protein expression and serine uptake, both of which can be re-stimulated with 100 nM estradiol (Figure 6D-E). Importantly, estrogen does not regulate serine uptake in ASCT2 KO cells (Figure 6F), indicating that estrogen-dependent control of serine uptake is through regulation of ASCT2 expression. To determine whether *SLC1A5* is a direct transcriptional target of ERα, we performed ChIP-qPCR using four distinct ERα antibodies and found that ERα binds near the *SLC1A5* transcription start site in an estrogen-dependent manner (Figure 6G), similar to the known ERα target *EGR3* (Figure S4B). Together, our results suggest that ERα regulates serine uptake by promoting transcription of its *bona fide* target gene *SLC1A5*.

**Figure 6.**
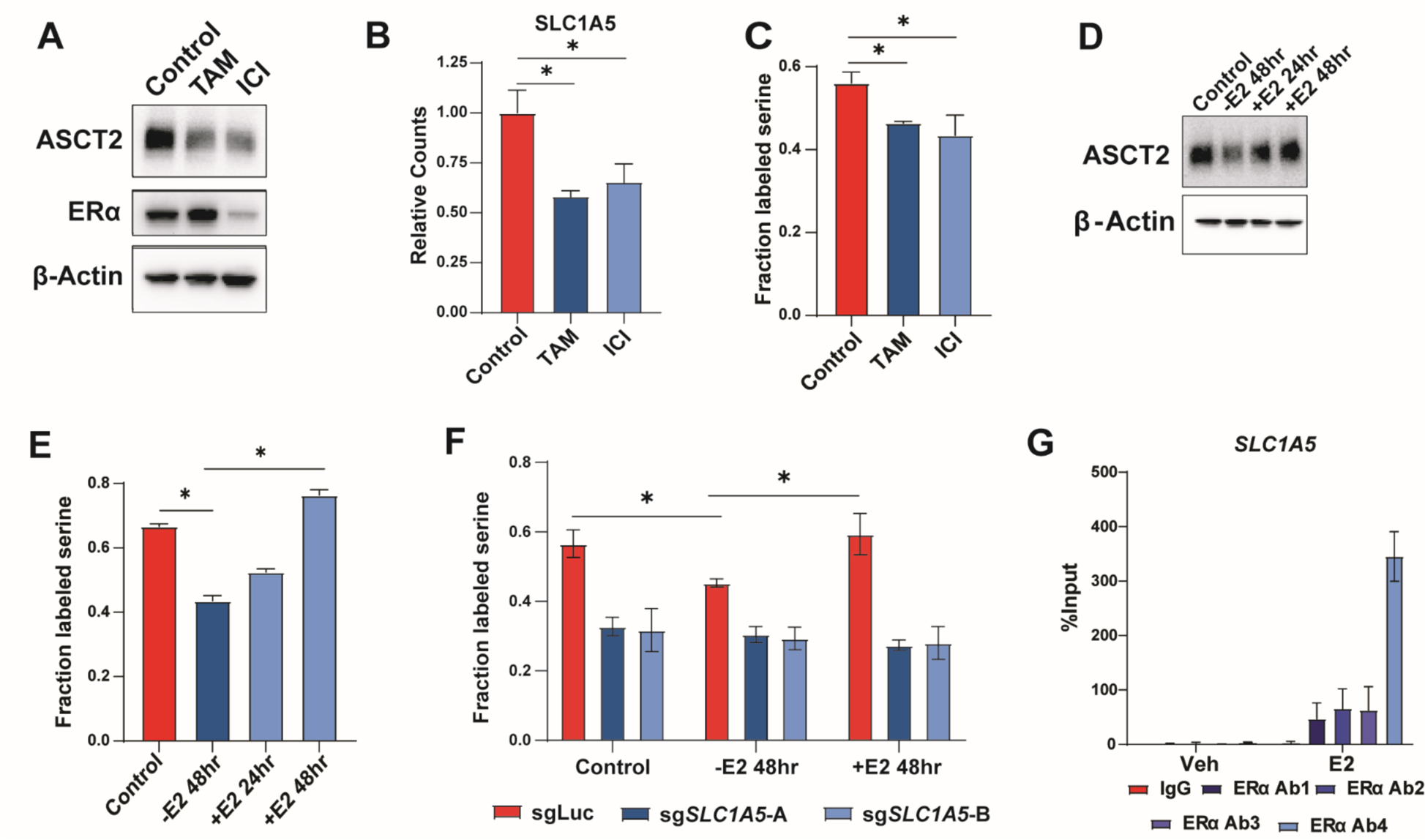
ERα promotes serine uptake via direct regulation of *SLC1A5*. A). Representative western blot of MCF7 cells cultured with DMSO (control), 1 µM tamoxifen (TAM), or 1 µM fulvestrant (ICI) for 48 hours. B). RNA-Seq data from MCF7 cells cultured with DMSO (Control), tamoxifen (TAM) or fulvestrant (ICI) for 48 hours. Values are the means ±SD of triplicate samples. *p < 0.05 by Welch’s t-test. C). Acute serine uptake in MCF7 cells treated with DMSO (Control), tamoxifen (TAM), or fulvestrant (ICI) for 48 hours. Values are the means ±SD of triplicate values from three independent experiments. *p < 0.05 by Welch’s t-test. D). Representative western blot of MCF7 cells starved of estrogen for 48 hours followed by add-back of 100nM estradiol for 24 or 48 hours. E). Acute serine uptake in MCF7 cells starved of estrogen for 48 hours followed by add-back of 100 nM estradiol for 24 or 48 hours prior to metabolite harvest. NS = No Starve, untreated cells. Values are the means ±SD of triplicate samples representative of two independent experiments. *p < 0.05 by Welch’s t-test. F). Acute serine uptake in MCF7 control (sgLuc) and *SLC1A5* KO (sg*SLC1A5*) cells under normal (NS) conditions, 48 hour estrogen starvation (-E2 48hr), and 48hr starve followed by 48 hour estradiol supplementation (+E2 48hr). Values are the means ±SD of triplicate samples representative of two independent experiments. *p < 0.05 by Welch’s t-test. G). ChIP-qPCR results from MCF7 cells treated with vehicle or 10 nM estradiol for 30 minutes. Four unique antibodies against ERα were used along with an IgG control antibody. Values are the means ±SD of triplicate samples from two independent qPCR experiments.

### Loss of ASCT2 inhibits tumor growth in combination with a serine-free diet

Traditional cell culture media contains supraphysiological concentrations of many amino acids^59^. Given that extracellular serine and glutamine levels impact serine uptake (Figure 4), we assessed the activity and importance of ASCT2 in more physiological conditions. First, we performed an additional CRISPRi screen assessing transporter essentiality in MCF7 cells growing in RPMI with amino acid concentrations similar to those found in human plasma (RPMI-PAA) and compared the magnitude of gene essentiality in RPMI-PAA to those found in RPMI (Figure 7A). The top hit in this screen was *SLC7A1*, a known arginine transporter^60^, that is likely more essential in RPMI-PAA due to the 90% lower arginine levels found in human plasma relative to RPMI^59^. Importantly, *SLC1A5* was the second strongest hit that was more essential in physiological amino acid levels than in RPMI (Figure 7A). We cultured control and ASCT2 KO cells in RPMI with plasma levels of serine (150 μM), glutamine (550 μM), or both, and observed increased dependence on ASCT2 in the more physiological conditions (Figure 7B), suggesting that lower serine and glutamine levels contribute to the increased *SLC1A5* essentiality in RPMI-PAA.

**Figure 7.**
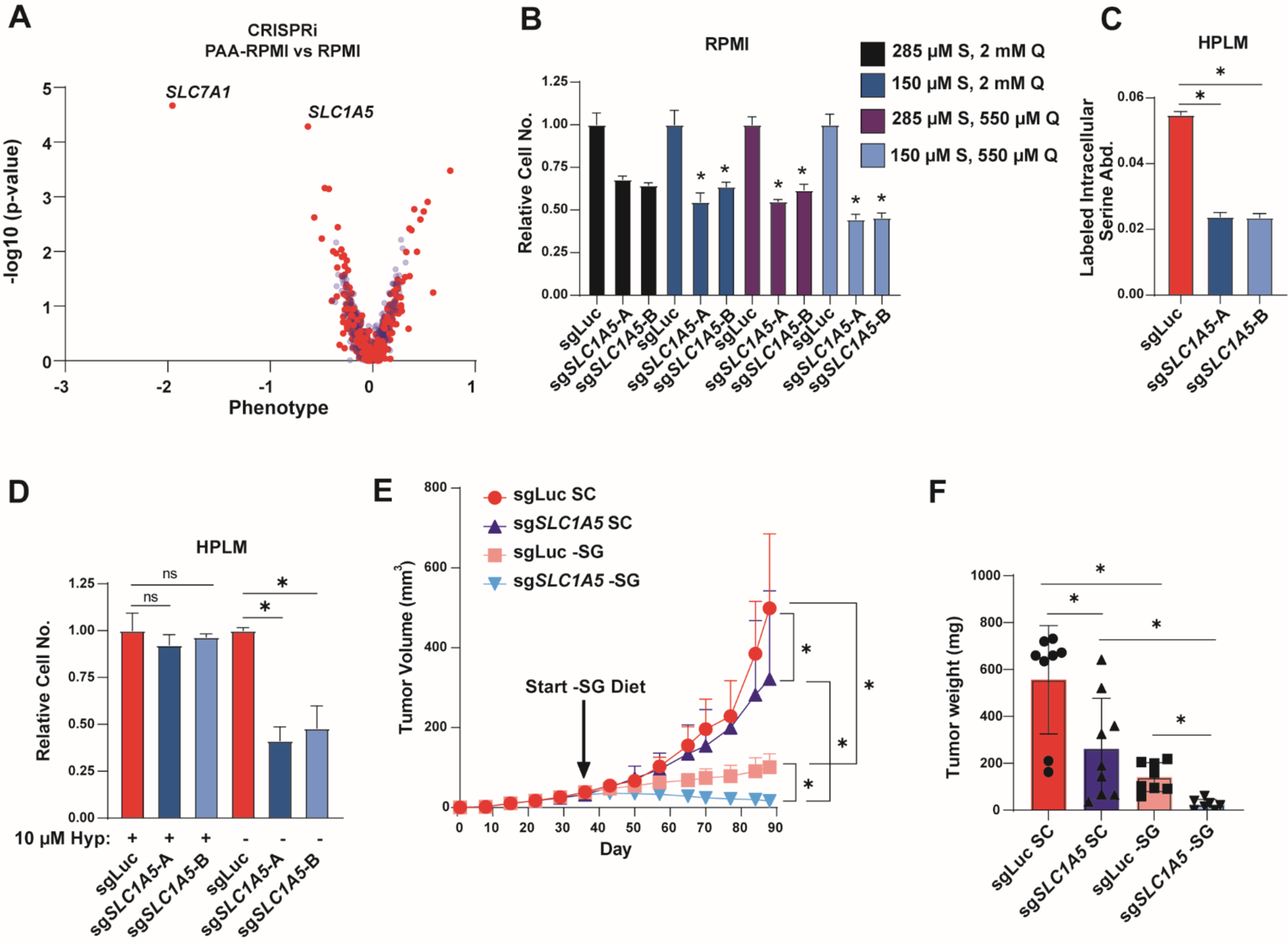
Loss of ASCT2 inhibits tumor growth in combination with serine-free diet. A). CRISPRi screen results from MCF7 cells grown in RPMI compared to RPMI-PAA (physiological amino acids). Blue are pseudogenes that represent the technical noise of the assay, red dots represent transporter genes. B). Growth assay from MCF7 control (sgLuc) and ASCT2 KO (sg*SLC1A5*) cells grown in RPMI with serine and glutamine concentrations at normal RPMI or physiological levels. Values are the means ±SD of triplicate samples representative of two independent experiments. *p < 0.05 by Welch’s t-test. C). Acute serine uptake in MCF7 control (sgLuc) and ASCT2 KO (sg*SLC1A5*) cells grown in HPLM. Values are the means ±SD of triplicate samples from an experiment representative of three independent experiments. *p < 0.05 by Welch’s t-test. D). Growth assay from MCF7 control (sgLuc) and ASCT2 KO (sg*SLC1A5*) cells grown in HPLM (150 μM serine) in the presence of absence of 10 µM hypoxanthine. Values are the means ±SD of triplicate samples from an experiment representative of two independent experiments. *p < 0.05 by Welch’s t-test. E). Tumor xenograft growth over time from MCF7 control (sgLuc) and ASCT2 KO (sg*SLC1A5*) xenografts. Mice were fed standard chow (SC) until tumors were established, then half of each cohort was switched to a serine and glycine free diet (-SG) for the remainder of the study. Values are the means ±SD of each group’s tumor volume measurements. *p < 0.05 by two-way ANOVA. F). Tumor weight at endpoint for each cohort of mice. *p < 0.05 by Welch’s t-test.

Next, we made use of human plasma-like media (HPLM), which (like RPMI-PAA) contains physiological concentrations of all amino acids, but also contains human plasma glucose and salt levels as well as 27 additional components found in human plasma but not RPMI or other traditional media^59^. Notably, loss of ASCT2 decreased serine uptake in HPLM to a similar degree as seen in RPMI medium (Figure 7C), suggesting that ASCT2 is still the primary serine transporter in physiological conditions. However, unlike in RPMI or RPMI-PAA, loss of ASCT2 does not strongly inhibit proliferation in plasma (150 µM) or low (50 µM) levels of serine in HPLM (Figure 7D). HPLM also contains 10 µM hypoxanthine, which we have previously shown to partially rescue the growth inhibitory effects of ASCT2 loss (Figure 5H). Indeed, removal of hypoxanthine from HPLM strongly reduces proliferation in ASCT2 KO cells (Figure 7D). This data suggests that there are aspects of more physiological conditions that increase (low serine/glutamine) and decrease (purine salvage substrates) the requirements for serine uptake mediated by ASCT2.

While HPLM can provide more physiological *in vitro* conditions, it cannot fully recapitulate the *in vivo* environment. As such, we grew control and ASCT2 KO MCF7 cells orthotopically in the mammary fat pad of nude mice and observed that loss of ASCT2 alone had a slight inhibitory effect on tumor growth, similar to our *in vitro* results (Figure 7E-F). To determine whether loss of ASCT2 sensitizes to a low serine environment *in vivo*, we switched a cohort of ASCT2 KO and control mice to a serine and glycine free diet (-SG) after tumors were established, where we observed the expected inhibitory effect of dietary serine starvation on control MCF7 tumors^20^. Critically, we found that dietary serine starvation in addition to ASCT2 KO not only inhibited tumor growth, but resulted in regression of almost all tumors (Figure 7E, 7F). These data suggest that although ASCT2 could be a viable therapeutic target on its own, the largest effects are likely to be seen when ASCT2 inhibition is combined with therapies that reduce serine availability *in vivo*.

## DISCUSSION

In the years since serine metabolism was first implicated in cancer^9,10^, considerable strides have been made towards understanding the central role serine plays in proliferating cells^61^. However, despite an appreciation for the importance of exogenous serine and the potential clinical relevance of dietary serine starvation in cancer treatment^17,62^, the transporters that mediate serine uptake in cancer cells have remained unidentified. In this manuscript we identify ASCT2 (*SLC1A5*) as the primary serine transporter in cancer cells. ASCT2 is well-known in the cancer metabolism field due to its contributions to glutamine uptake, but to our knowledge we are the first to demonstrate that ASCT2 is also an important contributor to serine uptake in cancer.

Because of its importance in glutamine metabolism, there have been considerable efforts to target ASCT2 in cancer^45,49,63^. Current ASCT2 inhibitors include the pseudo-metabolites benzylserine^64^ and GPNA^65^ as well as the small molecule V-9302^66^. While benzylserine and GPNA can reduce breast cancer cell growth by inhibiting uptake of several amino acids^64^, they will likely not be useful clinically due to the high millimolar concentrations required to be effective. Small molecule inhibitors like V-9302 have a higher potency and have been shown to significantly reduce glutamine uptake and cell growth in a number of cancer cell lines and xenograft models^66^, but V-9302 likely has low specificity for ASCT2 and its effects could be due to the inhibition of multiple transporters^67^. Nevertheless, efforts to specifically target ASCT2 are ongoing and will surely be aided by the recent publication of a complete human ASCT2 Cryo-EM structure^68,69^. Our discovery that ASCT2 is an important serine transporter could expand the potential utility of ASCT2 inhibitors to include treatment of serine auxotrophic tumors.

While our studies demonstrate that ASCT2 is the primary serine transporter in most cancer cells, it is apparent that there are likely one or more additional transporters that also contribute to serine uptake. Our targeted secondary transporter approach failed to identify strong contributions from ASCT1, SNAT1, SNAT2, LAT1, LAT2, or ATB0+ either alone or in combination with ASCT2 loss. Based on these results it is likely that there is a network of secondary serine transporters that all contribute to serine uptake, particularly when ASCT2 is absent. It is also possible that there are additional transporter(s) that have not yet been shown to transport serine *in vitro* that may also support serine uptake in cancer cells. Future work to understand this network of serine transporters will aid in the efforts to target serine metabolism more effectively.

Our *in vitro* and *in vivo* work both demonstrate that loss of ASCT2 has the greatest effect on cancer cell growth in low serine conditions. Importantly, unlike most previous reports showing that dietary serine starvation typically only slows tumor growth, we find that combined loss of ASCT2 with dietary serine starvation has the ability to induce regression of established xenograft tumors. This finding may be relevant clinically, as dietary serine starvation is currently being evaluated in a clinical trial for pancreatic cancers, a subset of which are auxotrophic for serine^23^. It is also important to note that there are environments within the body that are naturally low for serine, such as cerebrospinal fluid (CSF) and brain interstitial fluid, which contain some of the lowest levels of serine in the body^70–73^. While upregulation of serine biosynthesis promotes the outgrowth of triple-negative breast cancer brain metastases^74^, it is likely that luminal breast cancer brain metastases are even more dependent on ASCT2 in this naturally low serine environment. Therefore, further studies could identify scenarios where ASCT2 is a viable clinical target to treat serine auxotrophic tumors.

## METHODS

### CRISPRi and CRISPRa screens

We first generated and validated MCF7 cell lines that stably express either dCas9-KRAB (MCF7 CRISPRi) or dCas9-SunTag (MCF7 CRISPRa). MCF7 CRISPRi were prepared by transduction of MCF7 cells with lentiviral particles produced using vector pMH0001 (Addgene #85969) in the presence of 8 mg/mL polybrene (Sigma). A pure polyclonal population of dCas9-KRAB expressing cells was generated by two rounds of fluorescence activated cell sorting (FACS) gated on the top half of BFP^+^ cells (BD FACS Aria). The performance of MCF7 CRISPRi in knocking down endogenous genes was confirmed by individually targeting 3 control genes (ST3GAL4, SEL1L, DPH1) and measuring gene expression changes by RT-qPCR. To prepare MCF7 CRISPRa parental cells, MCF7 cells were first transduced with lentiviral particles produced using vector pHRdSV40-dCas9-10xGCN4_v4-P2A-BFP (Addgene #60903) in the presence of 8 mg/mL polybrene. BFP^+^ cells were isolated using one round of FACS and were subsequently transduced with lentiviral particles produced using vector pHRdSV40-scFv-GCN4-sfGFP-VP64-GB1-NLS (Addgene #60904). Cells with high GFP levels (top 25% of GFP^+^ cells) and high BFP levels (top 50% of BFP^+^ cells) were isolated by FACS and recovered in complete medium. Monoclonal cell lines were subsequently generated by limiting dilution of the sorted population in conditioned medium. Individual clones were tested for their ability to increase the expression of target control genes as previously reported^75^. A clone exhibiting robust growth and overexpression of target genes was selected as cell line MCF7 CRISPRa.

The design and construction of transporter CRISPRi and CRISPRa sgRNA libraries has been reported elsewhere^40^. The multiplicity of infection (MOI) of lentiviral supernatant produced from each library was determined by titration onto MCF7 CRISPRi and CRISPRa cell lines and quantification of the percentage of puromycin resistant cells. To prepare transporter MCF7 CRISPRi/a libraries, parental MCF7 CRISPRi/a cells were grown to a confluence of 80-90% in 2× 225 cm^2^ cell culture flask (Costar) in RPMI-1640 (Corning 10–040) supplemented with 10% (v/v) heat inactivated fetal bovine serum (FBS) (Gibco 10438026) and 2.5 g/L D-Glucose. Penicillin and streptomycin were added to all screen growth media to final concentrations of 100 U/mL and 100 μg/mL, respectively (Corning 30–002–CI). Cells (about 2× 30 million) were transduced by addition of library lentiviral supernatant in presence of 8 μg/mL polybrene to achieve an MOI of 0.3. 24 hr post-transduction, cells were harvested by trypsinization (trypsin-EDTA (0.05%), Gibco 25300-054) and resuspended in 4× 60 mL fresh medium in 4× 225 cm^2^ flasks. Starting 48 hr post-transduction, cells were cultured in complete medium + puromycin 1 μg/mL, and passaged as needed, for a total of 5 daily puromycin pulses. After recovery for 24 hr in puromycin-free RPMI-rich (see below), library cells were harvested and homogenized, and cell pellets were resuspended at 15-20 million/mL in phosphate-buffered saline (PBS, Corning 21–040 CV). 2× 12 million cells were washed 1× with PBS and cell pellets were stored at –80 °C (T = 0 samples). Screens were initiated by addition of 9 million cells (CRISPRi) or 7 million cells (CRISPRa) to 60 mL medium in 225 cm^2^ flasks.

To prepare media for screens, we first prepared complete RPMI lacking amino acids. 8.59 g of RPMI w/o amino acids, sodium phosphate (US Biological R8999–04A), 2.00 g sodium bicarbonate (Sigma S6014), and 0.80 g sodium phosphate dibasic (Sigma S0876) were diluted in 945 mL deionized H_2_O. After addition of 100 mL dialyzed fetal bovine serum (dFBS, Gibco, Thermo Fisher Scientific 26400-044) and 5 mL D-Glucose (500 g/L), the medium was sterilized via 0.22 μm membrane filtration. Amino acids (excluding serine and glycine) were added to this base medium from stock solutions to RPMI levels. To prepare low serine medium for CRISPRi and CRISPRa screens, serine was added to this medium to 50 μM and 19 μM, respectively, (instead of 285 μM in complete RPMI). These concentrations were determined in pre-screen titrations and chosen to achieve selective pressure favoring the isolation of hypersensitive hits in CRISPRi screens and resistant hits in CRISPRa screens. To make RPMI used as the control arm in low amino acid screens, serine and glycine were added to this medium to RPMI levels. To prepare complete RPMI medium with amino acids present at physiological levels screens (RPMI-PAA), base medium lacking amino acids was prepared as above, except that 965 mL deionized H_2_O was used. To that, amino acids were added to their concentration in human plasma using the same stock solutions as for RPMI above and as previously described^40^. All media were preheated and equilibrated to 37 °C/5% CO_2_ before use.

CRISPRi screens were conducted in RPMI low Serine, RPMI, and RPMI-PAA in a 225 cm^2^ flask each with 2 technical replicates. Libraries were maintained at a coverage of at least 1000 × at low cell density with frequent media changes and passaging as necessary for a total of 18 days. During that time, cells in RPMI, RPMI-PAA, and RPMI low Ser underwent 8.4, 7.8, and 6.4 population doublings, respectively. CRISPRa screens were conducted in RPMI and RPMI low Ser in a 225 cm^2^ flask each with 2 technical replicates over 20 days, and maintained as above. Cells in RPMI and RPMI low Ser underwent 6.8 and 1.3 population doublings, respectively.

At screen end, 6–12 million cells were collected by centrifugation, washed 1 × with PBS and stored at −80 °C. The abundance of each sgRNA in individual samples was determined as previously reported^40^. Briefly, genomic DNA (gDNA) was extracted from cell pellets using the QIAamp DNA Blood Mini Kit (Qiagen) and typical yields ranged from 40 to 80 μg gDNA. sgRNA barcodes were amplified using 23 cycles of PCR on the gDNA from at least 6 million cells as template, Q5 polymerase (NEB M0491L) and barcoded forward and reverse primers. Amplified PCR products (∼240–250 bp) were purified by agarose gel electrophoresis using the QIAquick gel extraction kit (Qiagen). Purified PCR products were quantified using the Qubit dsDNA high sensitivity assay kit (Thermo Fisher Scientific) and individual indexed libraries were mixed in equimolar ratio. Pooled libraries were sequenced on an Illumina NextSeq 500 platform using a 75 bp single read on a high output flow cell with a 2–5% PhiX spike-in. 12-20 million reads were obtained for each indexed sample. To determine counts per sgRNA sequence, reads were trimmed and aligned to the library of protospacers (93–94% of trimmed reads align). To estimate noise in the screens, simulated negative control genes (the same number as that of real transporter genes) were generated by randomly grouping 10 sgRNAs from the pool of 730 non-targeting control (NTC) sgRNAs present in the transporter libraries. For each gene (and simulated control gene), which is targeted by 10 sgRNAs, two metrics were calculated: (i) the mean of the strongest 7 rho phenotypes by absolute value (“Phenotype”), and (ii) the p-value of all 10 rho phenotypes compared to the 730 NTC sgRNAs (Mann-Whitney test). sgRNAs were required to have 100+ counts in at least one of the two conditions tested to be included in the analysis. To deal with noise associated with potential low count numbers, a pseudocount of 10 was added to all counts.

To determine the effect of transporter CRISPRi/a on serine transport, the two metrics were determined by comparison of sgRNA abundances in the RPMI low serine medium to those of RPMI. To determine differential transporter essentiality between RPMI-PAA and RPMI, the two metrics were determined by comparison of sgRNA abundances in RPMI-PAA to those in RPMI.

### Cell culture and media

MCF7, T47D, ZR751, HCC1806 and SUM149 cells were acquired from the Brugge Lab at Harvard Medical School. A498 ccRCC cells were acquired from Frank Mason at Vanderbilt University, and A549 and Calu6 lung cancer lines were acquired from Jiyeon Kim at the University of Illinois Chicago. Cell line identity was confirmed via STR analysis. All breast cell lines were of female origin. Cell lines were tested for mycoplasma using University of Illinois at Chicago Genome Research Core facilities. Unless otherwise noted, cells were grown in RPMI with 5% dialyzed FBS (Sigma) and Pen/Strep (Invitrogen) at 37 degrees C with 5% CO_2_. Human plasma like media (HPLM) was generated according to the published formulation^59^ with addition of 5% dialyzed FBS (Sigma) and Pen/Strep (Invitrogen). For RPMI containing altered serine or glutamine concentrations, media was formulated from RPMI powder lacking glucose and all amino acids (US Biological Life Sciences) with missing components added back as needed. Media were changed every other day for cell line propagation. For growth assays, cells were plated in complete RPMI, and medium was switched to low serine RPMI the next day and replenished every day for six days prior to counting with a Z1 Coulter Particle Counter (Beckman Coulter). For estrogen deprivation experiments, cells were plated in complete RPMI and allowed to attach overnight. The next day, cells were washed twice with PBS and fed RPMI lacking phenol red (Thermo Fisher Scientific) with 5% charcoal stripped FBS. Estradiol was added back to the media as needed at 100 nM. Tamoxifen (Sigma) and Fulvestrant (Sigma) were added to RPMI media at 1 µM for 48hours.

### Western blotting

Cells were lysed in RIPA buffer (Thermo Fisher) with protease and phosphatase inhibitors (Thermo Fisher) and 1 µM MG132 (Selleckchem). Protein concentration was determined by BCA assay (Thermo Fisher). When blotting for ASCT1, SNAT1, SNAT2, LAT2, ATB0+, lysates were subject to PNGase F treatment (New England Biolabs) to deglycosylate the transporter proteins. Briefly, 20-35 µg of protein was mixed with 1 µL of Glycoprotein Denaturing Buffer (10X) and water to reach a total volume of 10 µL. The reaction was heated to 100°C for 10 minutes. After heating, 2 µL of GlycoBuffer 2 (10X), 2 µL of 10% NP-40, 1 µL of PNGase F and water were added to a final volume of 20 µL. The samples were allowed to incubate at 37°C for one hour. Cell lysis samples that do not require PNGase F treatment were heated to 55°C prior to gel separation, as many transporter proteins are unstable at higher temperatures. Cell lysis protein samples were separated by gel electrophoresis on 4-20% ready-made Tris-Glycine gels (Invitrogen) and transferred at 110mV for one hour to PVDF membranes (Millipore). Membranes were blocked with 5% milk for 1 hour and incubated at 4°C overnight with one or more primary antibodies in 2% bovine serum albumin. Primary antibodies used were ASCT2 (Cell Signaling, 5245S), PHGDH (Sigma, HPA021241), PSAT1 (Thermo Fisher, PA5-22124), PSPH (Santa Cruz, sc-365183), ERα (Cell Signaling, 8644), ASCT1 (Santa Cruz, sc-393157), SNAT1 (Cell Signaling, 36057S), SNAT2 (Sigma, HPA035180), LAT1 (Cell Signaling, 5347S), LAT2 (OriGene, TA500513S), ATB0+ (US Biological, 41972), and β-Actin (Sigma, A1978). PVDF membranes were washed three times with tween 20-containing tris buffered saline before incubation with HRP-conjugated secondary antibodies (Bio-Rad). Images were detected using a ChemiDoc MP Imaging System (Bio-Rad).

### RT-qPCR

RNA was isolated with Trizol reagent (Thermo Fisher) and cDNA synthesis was performed using qScript cDNA synthesis Kits (QuantBio). RT-qPCR was run using SYBR Green on an ABI ViiA7 real-time PCR system (Applied Biosystems), and results were normalized to the expression of an *RPLP0* control primer set. The following primers were used: *SLC1A4* F *5’-GCTGTGGACTGGATTGTGG-3’, SLC1A4* R *5’- ATTCAGGTGGTGGAGAATGC-3’, SLC1A5* F *5’-CGGTCGACCATATCTCCTTG-3’*, *SLC1A5* R *5’- CTACATTGAGGACGGTACAGGA-3’*, *RPLPO* F *5’-ACGGGTACAAACGAGTCCTG-3’*, *RPLPO* R *5’- CGACTCTTCCTTGGCTTCAA-3’*.

### GC-MS metabolite analyses

For serine and glutamine acute uptake assays, cells were cultured in RPMI to 80% confluency, washed with PBS and fed RPMI containing U-^13^C_3_-^15^N-serine or U-^13^C_5_-^15^N_2_-glutamine (Cambridge Isotopes) for four (serine) or seven (glutamine) minutes. Labeled media was aspirated and cells were washed with saline and then lysed with ice cold GC-MS grade methanol (OmniSolv). Norvaline was added as an internal standard diluted in MilliQ water. GC-MS grade chloroform was added, and lysates were vortexed and centrifuged at 21,000 x g at 4 degrees C for 10 minutes. The polar metabolite fraction was separated and air dried prior to derivatization with MOX (Thermo Fisher, PI45950) and *N-tert-* butyldimethylchlorosilane (TBDMS) (Sigma-Aldrich). Where necessary, labelled intracellular serine and glutamine abundance were internally normalized to the average of over 20 additional analytes that were calculated by normalizing each analyte (from a library of 20+ analytes, excluding known ASCT2 substrates serine, glutamine, and threonine) to norvaline then dividing by the average of each analyte across all samples in the batch. Because manipulation of estrogen signaling affected the abundance of numerous analytes, these experiments were analyzed as the fraction of labeled Serine 390 M+4 out of the total pool of Serine 390.

For intracellular amino acid abundance determination after glutamine deprivation, cells were lysed as described and total abundance for each amino acid was normalized to the norvaline internal standard. All samples were analyzed by GC/MS using a HP-5MS Ultra Inert GC column (19091S-433UI, Agilent Technologies) installed in an Agilent 7890B gas chromatograph coupled to an Agilent 5779B mass spectrometer. Helium was used as the carrier gas. One microliter was injected (split inlet) at 280 degrees C. After injection, the GC oven was held at 60 degrees C for 1 minute before ramping to 320 degrees C at 10C/min and held for 9 minutes at the maximum temperature. The MS system operated under electron impact ionization mode at 70 eV and the MS source and quadrupole were held at 230 degrees C and 150 degrees C respectively. Peak areas were determined using MassHunter software.

### CRISPR knockout

Knockout of *SLC1A5*, *SLC1A4, SLC38A1, SLC38A2, SLC7A5, SLC7A8, SLC6A14,* and *PSAT1* were performed using lentiCRISPR v2 Puro or Hygro (Addgene, 52961 and 91977). The following oligos were cloned into BsmBI cut lentiCRISPR v2: *SLC1A5-A F: 5’-caccgAAGAGGTCCCAAAGGCAG-3’, SLC1A5-A R: 5’-aaacCTGCCTTTGGGACCTCTTc-3’, SLC1A5-B F: 5’-caccgTGCCCCACAGGAAGCGGT-3’, SLC1A5-B R: 5’-aaacACCGCTTCCTGTGGGGCAc-3’, SLC1A4-A F: 5’-caccgAGCAGGCGTGCCAGCTGG-3’, SLC1A4-A R: 5’-aaacCCAGCTGGCACGCCTGCTc-3’, SLC1A4-B F: 5’-caccgCTCCGTCCATGTTCACGG-3’, SLC1A4-B R: 5’-aaacCCGTGAACATGGACGGAGc-3’, SLC38A1-A F: 5’-caccgATGGTGTATGAAAAGCTG-3’, SLC38A1-A R: 5’-aaacCAGCTTTTCATACACCATc-3’, SLC38A1-B F: 5’-caccgTTCTTCAAGAGACACAG-3’, SLC38A1-B R: 5’-aaacCTGTGTCTCTTGAAGAAc-3’, SLC38A2-A F: 5’-caccgATATTTGGGATATACCAG-3’, SLC38A2-A R: 5’-aaacCTGGTATATCCCAAATATc-3’, SLC38A2-B F: 5’-caccgAGCAGCTTCCACAGGACA-3’, SLC38A2-B R: 5’-aaacTGTCCTGTGGAAGCTGCTc-3’, SLC7A5-A F: 5’-caccgACGACAGCATCTGCTCGG-3’, SLC7A5-A R: 5’-aaacCCGAGCAGATGCTGTCGTc-3’, SLC7A5-B F: 5’-caccgTGTGGGTGGATCATGGAG-3’, SLC7A5-B R: 5’-aaacCTCCATGATCCACCCACAc-3’, SLC7A8-A F: 5’-caccgCATCCAACGCCGTCGCTG-3’*, *SLC7A8-A R: 5’-aaacCAGCGACGGCGTTGGATGc-3’, SLC7A8-B F: 5’-caccgTCAGGCTTCTTCCAGCGA-3’, SLC7A8-B R: 5’-aaacTCGCTGGAAGAAGCCTGAc-3’, SLC6A14-A F: 5’-caccgACACTCCAGAAAGAACAA-3’, SLC6A14-A R: 5’-aaacTTGTTCTTTCTGGAGTGTc-3’, SLC6A14-B F: 5’-caccgATATCTGACCTACAGCAA-3’, SLC6A14-B R: 5’-aaacTTGCTGTAGGTCAGATATc-3’, PSAT1 F: 5’-caccgACCGAGGGGCACTCTCGG-3’, PSAT1 R: 5’-aaacCCGAGAGTGCCCCTCGGTc-3’*

### Over-expression studies

For *SLC1A5* and *PSAT1* over expression, human *SLC1A5* and *PSAT1* were cloned into pLenti CMV Puro and Neo DEST, respectively (Addgene, 17452, 17392), using the gateway cloning system (Thermo Fisher). To generate lentiviral particles, HEK293T cells were transduced using PAX2, VSVG, polyethylenimine (Polysciences) and the lentiviral plasmid of interest. Viral supernatants were collected on days 2, 3, and 4 after transduction. After infection with polybrene (Sigma), cells were drug selected until mock infected cells were completely cleared, after which antibiotic was removed for further propagation.

### ChIP-qPCR

Chromatin immunoprecipitation was performed as previously described^76^. MCF7 cells underwent crosslinking in 1% formaldehyde in PBS. Precipitations utilized protein A Dynabeads (10003D, Invitrogen) coated with one of four ERα antibodies (Milipore 06-935, Abcam Ab3575, Cell Signaling #8644, Santa Cruz HC20). Excess antibody was washed before beginning pulldown. Samples underwent pulldown at 4°C while rotating for 16 hours, after which beads were washed followed by treatment with elution buffer (0.1 M NaHCO_3_, 1% SDS) and de-crosslinking overnight at 65°C. DNA was purified using a QIAquick PCR Purification Kit followed by qPCR as described above. The following primers were used: *SLC1A5* F: *5’-TGCTAGCCCTGAGGCATTGT*-3’, *SLC1A5* R: 5’- *ATGCAAGCTGTCCAGGGTATT*-3’.

### Purine biosynthesis tracing

MCF7 control (sgLuc) and AST2 KO (sg*SLC1A5*) cells were plated in triplicate and allowed to attach overnight. The next day, cells were washed with PBS and fed U-^13^C-serine at complete RPMI (285 µM) or low (50 µM) concentrations for eight hours. After incubation with labeled serine media, we collected 1mL of media from each well to be stored at −20°C and the remaining media was discarded. Cells were washed with 1mL of ice cold PBS on ice followed by 500 µL of 80% LC/MS-grade methanol (OmniSolv). The cells were then incubated at −80°C for 30 minutes. Each well was scraped, and the lysate-methanol mixture was transferred to 1.5mL Eppendorf tubes, which were centrifuged at 17,000 x *g* for 20 minutes at 4°C. Supernatant was collected from each tube and stored at −80°C until LC/MS analysis.

U-^13^C_3_-Serine labeled samples were analyzed using the Vanquish UPLC system coupled to a Q Exactive HF (QE-HF) mass spectrometer equipped with HESI (Thermo Fisher). The instrument conditions were optimized based on established methods^77^. 5 µL of all samples were injected by an auto-sampler and separated by an Atlantis Premier BEH Z-HILIC VanGuard FIT column (2.1 mm x 150 mm, 2.5 µm (Waters)) using a gradient elution of water containing 10mM ammonium carbonate and 0.05% ammonium hydroxide (Solvent A) and acetonitrile (Solvent B) at a flow rate of 0.25 mL/min. Injected samples were eluted by linear gradient from 80% to 20% Solvent B in 13 minutes. This percentage was maintained for 2 minutes and equilibrium time of 4.9 minutes. The eluted metabolites were detected using negative electrospray ionization mode. Full MS spectra were captured from 65 to 975 m/z at 120,000 resolutions with an automatic gain control (AGC) set to 3×10^6^ charges. The capillary temperature and voltage were 320°C and 3.5kV, respectively.

Following the initial conversion of the .raw data files to the .cdf format using Xcalibur (Version 4.0), we performed further data processing to facilitate targeted metabolomics analysis. This subsequent processing was carried out utilizing El-Maven (Version 0.10.0 or 0.12.0), with default parameters maintained except: the ionization mode was set to negative, an isotopic tracer of C13 was employed, and the extracted-ion chromatogram (EIC) extraction window was established at ±15.00 ppm. Metabolite identification relied on matching retention time and precursor ion m/z with our established library^78^. Peak intensity in EIC was measured using AreaTop. Isotope correction utilized El-Maven’s output in IsoCor (Version 1.0 or 2.2.0), with parameters: 13C tracer, ‘Low resolution’, ‘Correct natural abundance’ selected; isotopic purity: 12C (0.01), 13C (0.99).

### Mouse studies

All experiments using mice were approved by the University of Illinois at Chicago Animal Care Committee. MCF7 (5 x 10^6^) sgLuc and sg*SLC1A5*-B cells were injected into the mammary fat pad at the #4 and #9 mammary glands of 6 to 8-week old athymic nude-foxn1^nu^ female mice (Inotiv). Cell suspensions were injected at a volume of 50 µL in growth factor reduced Matrigel (Corning). Estrogen (E2) was added to the drinking water in each cage and replaced every 3-4 days. Tumor growth was measured over time via caliper measurements and tumor volume was calculated with the formula: volume = ½ (width^2^ x length). Two cohorts of mice (one sgLuc (N=5) and one sg*SLC1A5*-B (N=5)) were switched from standard chow to a serine and glycine free diet (Envigo, TD.160752) after palpable tumors had formed and food was replenished twice per week. At endpoint, mice were euthanized according to institutional guidelines.

### RNA-Sequencing

MCF7 cells were treated with DMSO vehicle control, Tamoxifen (1 µM), or Fulvestrant (1 µM) in HPLM with or without serine and glycine (SG), with 5% dialyzed FBS (Sigma) and Pen/Strep (Invitrogen). After 48 hours of treatment, RNA was isolated from the cells using Trizol reagent (Thermo Fisher). RNA samples were sent to Novogene for human mRNA sequencing using Illumina NovaSeq 6000 platform and paired end 150 bp (PE150) sequencing strategy. This service included library preparation prior to sequencing, and quantification analysis, including mapping reads to reference genome, gene expression quantification, differential expression analysis, pathway analysis.

### Charcoal Stripped Serum

A charcoal-dextran solution (5 mg of charcoal (Fisher), 500 mg dextran (Sigma), 50 mL 0.15 M NaCl (Fisher) was added to a 500 mL bottle of FBS (Sigma) and the mixture was incubated for 45 minutes at 56°C with 180 rpm rotation. After shaking, the mixture was centrifuged at 4000 rpm for 20 minutes at room temperature. An additional 25 mL of charcoal-dextran solution was added and the same rotation/incubation and centrifugation was performed two additional times. The mixture was then sterile-filtered through a 0.45 µm bottle-top filter and the filtered again through a 0.22 µm bottle-top filter. Charcoal-stripped FBS was aliquoted and stored at −20°C.

### Statistical Analysis

The transporter-specific CRISPR screen compares the individual transporter gene average of the strongest 7 phenotype scores from the 10 sgRNAs used for each gene with the p-value of all 10 phenotypes compared to the 730 non-targeting control gene sgRNAs (Mann-Whitney test). The results displayed as volcano plots represent a single score calculated by multiplying the phenotype score with the −log10(p-value). Only sgRNAs with a count greater than 100 in at least one of the media conditions were included in the final analysis. Volcano plots were generated in GraphPad Prism. All other statistical tests (as indicated in the figure legends) were performed using GraphPad Prism.

## Acknowledgements

We thank Dr. Jiyeon Kim and Dr. Isaac Harris for their helpful discussions and reagents. This work was supported by National Cancer Institute R37 CA251216 J.L.C, U54-CA225088 P.K.S, and The Harvard Ludwig Center for Cancer Research and the Termeer Foundation P.K.S

## Author contributions

J.L.C and K.O.C conceptualization, J.L.C, K.O.C, C.C, M.E.O, H.Z, Y.K, S.E.S, P.B, V.R, R.S, I.B, J.F, P.K.S, G.M.D methodology, J.L.C and K.O.C formalization, J.L.C, K.O.C, C.C, M.E.O, H.Z, Y.K, S.E.S, P.B, V.R, R.S investigation, J.L.C, R.S, I.B, J.F, P.K.S, G.M.D resources, J.L.C and K.O.C writing original draft, J.L.C, K.O.C, C.C, Y.K, P.B, R.S, I.B, J.F, P.K.S, G.M.D review and editing, J.L.C and K.O.C visualization, J.L.C, I.B, J.F, P.K.S, G.M.D supervision, J.L.C project admin, J.L.C. funding acquisition

---

**Figure S1:**
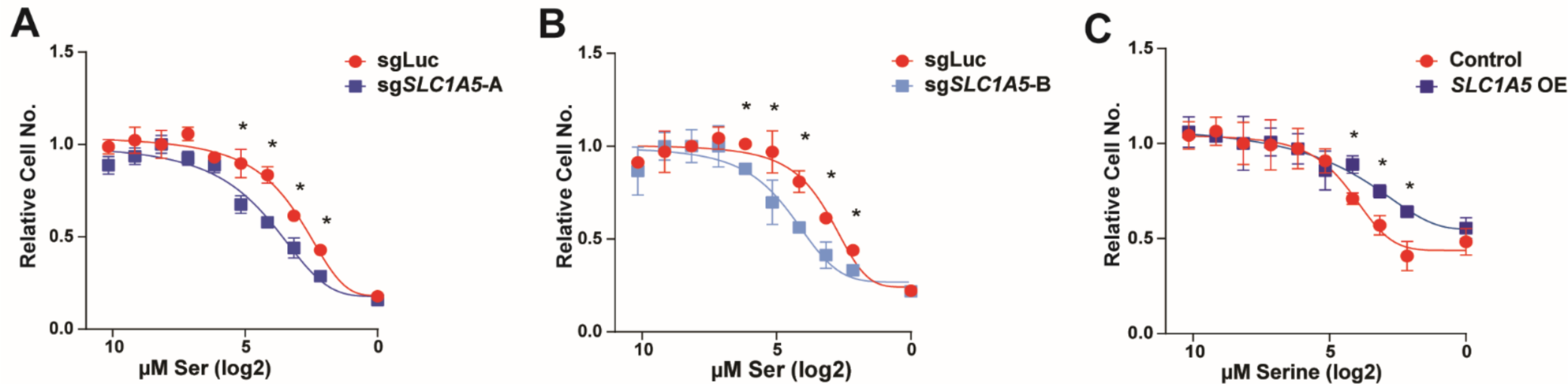
Identification of ASCT2 as a serine transporter in luminal breast cancer cells. A-B) Serine dose curves with MCF7 control (sgLuc) and ASCT2 KO (sg*SLC1A5*) cells. Values are the means ±SD of triplicate samples from a representative experiment of 3 independent experiments. *p < 0.05 by Welch’s t-test. Cell numbers normalized to each group’s count at normal RPMI serine dose. C). Serine dose growth curve with MCF7 control and ASCT2 over-expressing (*SLC1A5* OE) cells. Values are the means ±SD of triplicate samples from an experiment representative of 3 independent experiments. *p < 0.05 by Welch’s t-test comparing individual serine doses. Cell numbers normalized to each group’s count at normal RPMI serine dose.

**Figure S2.**
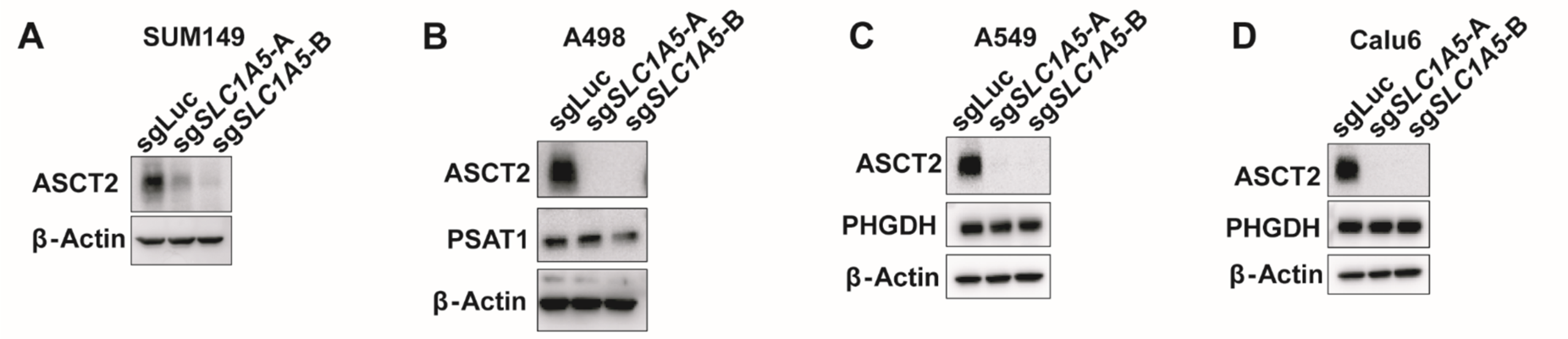
ASCT2 contributes to serine uptake in non-auxotrophic cells. A-D). Representative western blot from SUM149 (A), A498 (B), A549 (C), and Calu6 (D) control (sgLuc) and ASCT2 KO (sg*SLC1A5*) cells.

**Figure S3.**
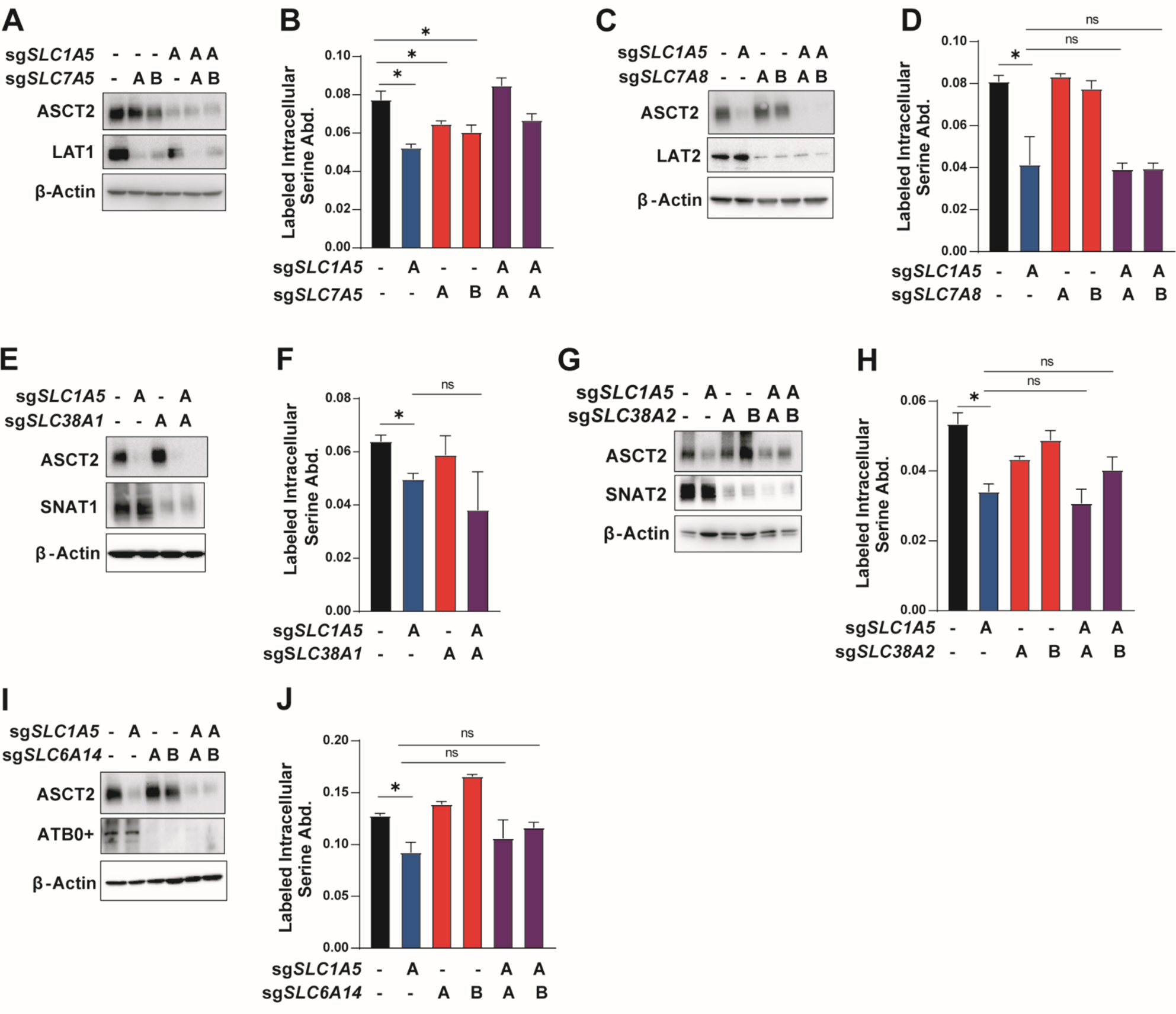
Transporter compensation in the absence of ASCT2. A). Western blot from single and double LAT1 (sg*SLC7A5*) and ASCT2 (sg*SLC1A5*) KO MCF7 cells. B). Acute serine uptake for single and double LAT1 (sg*SLC7A5*) and ASCT2 (sg*SLC1A5*) KO MCF7 cells. Values are the mean ±SD of triplicate samples from an experiment representative of 2 independent experiments. *p < 0.05 by Welch’s t-test. C). Western bot from single and double LAT2 (sg*SLC7A8*) and ASCT2 (sg*SLC1A5*) KO MCF7 cells. D). Acute serine uptake for single and double LAT2 (sg*SLC7A8*) and ASCT2 (sg*SLC1A5*) KO MCF7 cells. Values are the mean ±SD of triplicate samples from an experiment representative of 2 independent experiments. *p < 0.05 by Welch’s t-test. E). Western blot from single and double SNAT1 (sg*SLC38A1*) and ASCT2 (sg*SLC1A5*) KO MCF7 cells. F). Acute serine uptake for single and double SNAT1 (sg*SLC38A1*) and ASCT2 (sg*SLC1A5*) KO MCF7 cells. Values are the mean ±SD of triplicate samples from an experiment representative of 3 independent experiments. *p < 0.05 by Welch’s t-test. G). Western blot for single and double SNAT2 (sg*SLC38A2*) and ASCT2 (sg*SLC1A5*) KO MCF7 cells. H). Acute serine uptake for single and double SNAT2 (sg*SLC38A2*) and ASCT2 (sg*SLC1A5*) KO MCF7 cells. Values are the mean ±SD of triplicate samples from an experiment representative of 3 independent experiments. *p < 0.05 by Welch’s t-test. I). Western blot for single and double ATB0+ (sg*SLC6A14*) and ASCT2 (sg*SLC1A5*) KO MCF7 cells. J). Acute serine uptake for single and double ATB0+ (sg*SLC6A14*) and ASCT2 (sg*SLC1A5*) KO MCF7 cells. Values are the mean ±SD of triplicate samples from an experiment representative of 3 independent experiments. *p < 0.05 by Welch’s t-test.

**Figure S4.**
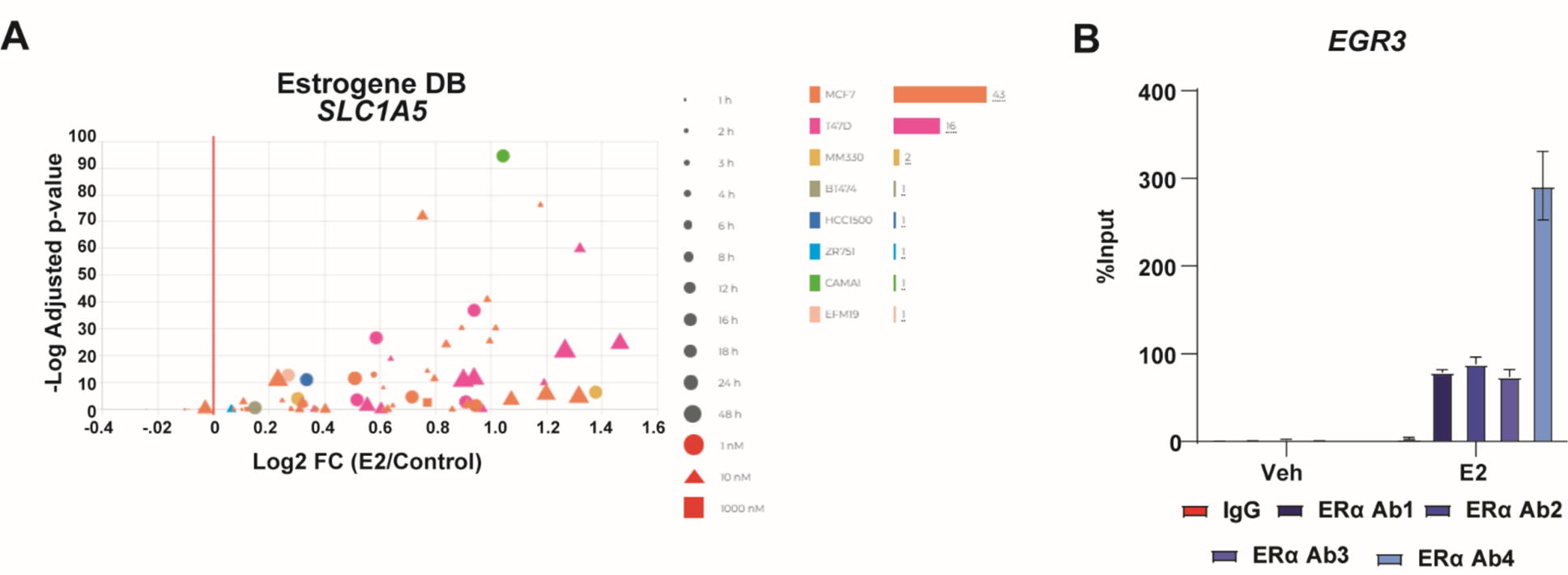
ERα promotes serine uptake via direct regulation of *SLC1A5*. A). RNA-Seq Analysis of *SLC1A5* expression after stimulation with different doses of estradiol in breast cancer cell lines from Estrogene DB (www.estrogene.org). B). ChIP-qPCR results from MCF7 cells treated with vehicle or 10 nM estradiol for 30 minutes. Four unique antibodies against ERα were used along with an IgG control antibody. Values are the means ±SD of triplicate samples from two independent qPCR experiments. *EGR3* is a known transcriptional target of ERα and therefore acts as a positive control.

## References

1. Vettore, L., Westbrook, R.L., and Tennant, D.A. (2020). New aspects of amino acid metabolism in cancer. Br. J. Cancer 122, 150–156. 10.1038/s41416-019-0620-5.

2. Hosios, A.M., Hecht, V.C., Danai, L.V., Johnson, M.O., Rathmell, J.C., Steinhauser, M.L., Manalis, S.R., and Vander Heiden, M.G. (2016). Amino Acids Rather than Glucose Account for the Majority of Cell Mass in Proliferating Mammalian Cells. Dev. Cell 36, 540–549. 10.1016/j.devcel.2016.02.012.

3. Geck, R.C., and Toker, A. (2016). Nonessential amino acid metabolism in breast cancer. Adv. Biol. Regul. 62, 11–17. 10.1016/j.jbior.2016.01.001.

4. Fan, T.W.M., Bruntz, R.C., Yang, Y., Song, H., Chernyavskaya, Y., Deng, P., Zhang, Y., Shah, P.P., Beverly, L.J., Qi, Z., et al. (2019). De novo synthesis of serine and glycine fuels purine nucleotide biosynthesis in human lung cancer tissues. J. Biol. Chem. 294, 13464–13477. 10.1074/jbc.RA119.008743.

5. Maddocks, O.D.K., Labuschagne, C.F., Adams, P.D., and Vousden, K.H. (2016). Serine Metabolism Supports the Methionine Cycle and DNA/RNA Methylation through De Novo ATP Synthesis in Cancer Cells. Mol. Cell 61, 210–221. 10.1016/j.molcel.2015.12.014.

6. Reid, M.A., Allen, A.E., Liu, S., Liberti, M.V., Liu, P., Liu, X., Dai, Z., Gao, X., Wang, Q., Liu, Y., et al. (2018). Serine synthesis through PHGDH coordinates nucleotide levels by maintaining central carbon metabolism. Nat. Commun. 9, 5442. 10.1038/s41467-018-07868-6.

7. Gao, X., Lee, K., Reid, M.A., Sanderson, S.M., Qiu, C., Li, S., Liu, J., and Locasale, J.W. (2018). Serine Availability Influences Mitochondrial Dynamics and Function through Lipid Metabolism. Cell Rep. 22, 3507–3520. 10.1016/j.celrep.2018.03.017.

8. Muthusamy, T., Cordes, T., Handzlik, M.K., You, L., Lim, E.W., Gengatharan, J., Pinto, A.F.M., Badur, M.G., Kolar, M.J., Wallace, M., et al. (2020). Serine restriction alters sphingolipid diversity to constrain tumour growth. Nature 586, 790–795. 10.1038/s41586-020-2609-x.

9. Possemato, R., Marks, K.M., Shaul, Y.D., Pacold, M.E., Kim, D., Birsoy, K., Sethumadhavan, S., Woo, H.-K., Jang, H.G., Jha, A.K., et al. (2011). Functional genomics reveal that the serine synthesis pathway is essential in breast cancer. Nature 476, 346–350. 10.1038/nature10350.

10. Locasale, J.W., Grassian, A.R., Melman, T., Lyssiotis, C.A., Mattaini, K.R., Bass, A.J., Heffron, G., Metallo, C.M., Muranen, T., Sharfi, H., et al. (2011). Phosphoglycerate dehydrogenase diverts glycolytic flux and contributes to oncogenesis. Nat. Genet. 43, 869–874. 10.1038/ng.890.

11. Pacold, M.E., Brimacombe, K.R., Chan, S.H., Rohde, J.M., Lewis, C.A., Swier, L.J.Y.M., Possemato, R., Chen, W.W., Sullivan, L.B., Fiske, B.P., et al. (2016). A PHGDH inhibitor reveals coordination of serine synthesis and one-carbon unit fate. Nat. Chem. Biol. 12, 452–458. 10.1038/nchembio.2070.

12. Mullarky, E., Lucki, N.C., Beheshti Zavareh, R., Anglin, J.L., Gomes, A.P., Nicolay, B.N., Wong, J.C.Y., Christen, S., Takahashi, H., Singh, P.K., et al. (2016). Identification of a small molecule inhibitor of 3-phosphoglycerate dehydrogenase to target serine biosynthesis in cancers. Proc. Natl. Acad. Sci. 113, 1778–1783. 10.1073/pnas.1521548113.

13. Labuschagne, C.F., van den Broek, N.J.F., Mackay, G.M., Vousden, K.H., and Maddocks, O.D.K. (2014). Serine, but Not Glycine, Supports One-Carbon Metabolism and Proliferation of Cancer Cells. Cell Rep. 7, 1248–1258. 10.1016/j.celrep.2014.04.045.

14. Amelio, I., Melino, G., and Frezza, C. (2017). Exploiting tumour addiction with a serine and glycine-free diet. Cell Death Differ. 24, 1311–1313. 10.1038/cdd.2017.83.

15. Li, X., Gracilla, D., Cai, L., Zhang, M., Yu, X., Chen, X., Zhang, J., Long, X., Ding, H.-F., and Yan, C. (2021). ATF3 promotes the serine synthesis pathway and tumor growth under dietary serine restriction. Cell Rep. 36, 109706. 10.1016/j.celrep.2021.109706.

16. Gravel, S.-P., Hulea, L., Toban, N., Birman, E., Blouin, M.-J., Zakikhani, M., Zhao, Y., Topisirovic, I., St-Pierre, J., and Pollak, M. (2014). Serine Deprivation Enhances Antineoplastic Activity of Biguanides. Cancer Res. 74, 7521–7533. 10.1158/0008-5472.CAN-14-2643-T.

17. Tajan, M., Hennequart, M., Cheung, E.C., Zani, F., Hock, A.K., Legrave, N., Maddocks, O.D.K., Ridgway, R.A., Athineos, D., Suárez-Bonnet, A., et al. (2021). Serine synthesis pathway inhibition cooperates with dietary serine and glycine limitation for cancer therapy. Nat. Commun. 12, 366. 10.1038/s41467-020-20223-y.

18. Falcone, M., Uribe, A.H., Papalazarou, V., Newman, A.C., Athineos, D., Stevenson, K., Sauvé, C.-E.G., Gao, Y., Kim, J.K., Del Latto, M., et al. (2022). Sensitisation of cancer cells to radiotherapy by serine and glycine starvation. Br. J. Cancer 127, 1773–1786. 10.1038/s41416-022-01965-6.

19. Maddocks, O.D.K., Berkers, C.R., Mason, S.M., Zheng, L., Blyth, K., Gottlieb, E., and Vousden, K.H. (2013). Serine starvation induces stress and p53-dependent metabolic remodelling in cancer cells. Nature 493, 542–546. 10.1038/nature11743.

20. Choi, B.-H., Rawat, V., Högström, J., Burns, P.A., Conger, K.O., Ozgurses, M.E., Patel, J.M., Mehta, T.S., Warren, A., Selfors, L.M., et al. (2022). Lineage-specific silencing of PSAT1 induces serine auxotrophy and sensitivity to dietary serine starvation in luminal breast tumors. Cell Rep. 38, 110278. 10.1016/j.celrep.2021.110278.

21. Dalton, W.B., Helmenstine, E., Walsh, N., Gondek, L.P., Kelkar, D.S., Read, A., Natrajan, R., Christenson, E.S., Roman, B., Das, S., et al. Hotspot SF3B1 mutations induce metabolic reprogramming and vulnerability to serine deprivation. J. Clin. Invest. 129, 4708–4723. 10.1172/JCI125022.

22. Baksh, S.C., Todorova, P.K., Gur-Cohen, S., Hurwitz, B., Ge, Y., Novak, J.S.S., Tierney, M.T., dela Cruz-Racelis, J., Fuchs, E., and Finley, L.W.S. (2020). Extracellular serine controls epidermal stem cell fate and tumour initiation. Nat. Cell Biol., 1–12. 10.1038/s41556-020-0525-9.

23. Banh, R.S., Biancur, D.E., Yamamoto, K., Sohn, A.S.W., Walters, B., Kuljanin, M., Gikandi, A., Wang, H., Mancias, J.D., Schneider, R.J., et al. (2020). Neurons Release Serine to Support mRNA Translation in Pancreatic Cancer. Cell 183, 1202–1218.e25. 10.1016/j.cell.2020.10.016.

24. Scerri, T.S., Quaglieri, A., Cai, C., Zernant, J., Matsunami, N., Baird, L., Scheppke, L., Bonelli, R., Yannuzzi, L.A., Friedlander, M., et al. (2017). Genome-wide analyses identify common variants associated with macular telangiectasia type 2. Nat. Genet. 49, 559–567. 10.1038/ng.3799.

25. Douce, J.L., Maugard, M., Veran, J., Matos, M., Jégo, P., Vigneron, P.-A., Faivre, E., Toussay, X., Vandenberghe, M., Balbastre, Y., et al. (2020). Impairment of Glycolysis-Derived l-Serine Production in Astrocytes Contributes to Cognitive Deficits in Alzheimer’s Disease. Cell Metab. 31, 503–517.e8. 10.1016/j.cmet.2020.02.004.

26. Handzlik, M.K., Gengatharan, J.M., Frizzi, K.E., McGregor, G.H., Martino, C., Rahman, G., Gonzalez, A., Moreno, A.M., Green, C.R., Guernsey, L.S., et al. (2023). Insulin-regulated serine and lipid metabolism drive peripheral neuropathy. Nature 614, 118–124. 10.1038/s41586-022-05637-6.

27. Gantner, M.L., Eade, K., Wallace, M., Handzlik, M.K., Fallon, R., Trombley, J., Bonelli, R., Giles, S., Harkins-Perry, S., Heeren, T.F.C., et al. (2019). Serine and Lipid Metabolism in Macular Disease and Peripheral Neuropathy. N. Engl. J. Med. 381, 1422–1433. 10.1056/NEJMoa1815111.

28. Eade, K., Gantner, M.L., Hostyk, J.A., Nagasaki, T., Giles, S., Fallon, R., Harkins-Perry, S., Baldini, M., Lim, E.W., Scheppke, L., et al. (2021). Serine biosynthesis defect due to haploinsufficiency of PHGDH causes retinal disease. Nat. Metab. 3, 366–377. 10.1038/s42255-021-00361-3.

29. Bonelli, R., Woods, S.M., Ansell, B.R.E., Heeren, T.F.C., Egan, C.A., Khan, K.N., Guymer, R., Trombley, J., Friedlander, M., Bahlo, M., et al. (2020). Systemic lipid dysregulation is a risk factor for macular neurodegenerative disease. Sci. Rep. 10, 12165. 10.1038/s41598-020-69164-y.

30. Oppedisano, F., Pochini, L., Galluccio, M., Cavarelli, M., and Indiveri, C. (2004). Reconstitution into liposomes of the glutamine/amino acid transporter from renal cell plasma membrane: functional characterization, kinetics and activation by nucleotides. Biochim. Biophys. Acta BBA - Biomembr. 1667, 122–131. 10.1016/j.bbamem.2004.09.007.

31. Segawa, H., Fukasawa, Y., Miyamoto, K., Takeda, E., Endou, H., and Kanai, Y. (1999). Identification and Functional Characterization of a Na+-independent Neutral Amino Acid Transporter with Broad Substrate Selectivity*. J. Biol. Chem. 274, 19745–19751. 10.1074/jbc.274.28.19745.

32. Kanai, Y., Segawa, H., Miyamoto, K., Uchino, H., Takeda, E., and Endou, H. (1998). Expression Cloning and Characterization of a Transporter for Large Neutral Amino Acids Activated by the Heavy Chain of 4F2 Antigen (CD98)*. J. Biol. Chem. 273, 23629–23632. 10.1074/jbc.273.37.23629.

33. Arriza, J.L., Kavanaugh, M.P., Fairman, W.A., Wu, Y.N., Murdoch, G.H., North, R.A., and Amara, S.G. (1993). Cloning and expression of a human neutral amino acid transporter with structural similarity to the glutamate transporter gene family. J. Biol. Chem. 268, 15329–15332. 10.1016/S0021-9258(18)82257-8.

34. Mackenzie, B., Schäfer, M.K.-H., Erickson, J.D., Hediger, M.A., Weihe, E., and Varoqui, H. (2003). Functional Properties and Cellular Distribution of the System A Glutamine Transporter SNAT1 Support Specialized Roles in Central Neurons*. J. Biol. Chem. 278, 23720–23730. 10.1074/jbc.M212718200.

35. Kaplan, E., Zubedat, S., Radzishevsky, I., Valenta, A.C., Rechnitz, O., Sason, H., Sajrawi, C., Bodner, O., Konno, K., Esaki, K., et al. (2018). ASCT1 (Slc1a4) transporter is a physiologic regulator of brain D - serine and neurodevelopment. Proc. Natl. Acad. Sci. 115, 9628–9633. 10.1073/pnas.1722677115.

36. Damseh, N., Simonin, A., Jalas, C., Picoraro, J.A., Shaag, A., Cho, M.T., Yaacov, B., Neidich, J., Al-Ashhab, M., Juusola, J., et al. (2015). Mutations in SLC1A4, encoding the brain serine transporter, are associated with developmental delay, microcephaly and hypomyelination. J. Med. Genet. 52, 541–547. 10.1136/jmedgenet-2015-103104.

37. Rosenberg, D., Artoul, S., Segal, A.C., Kolodney, G., Radzishevsky, I., Dikopoltsev, E., Foltyn, V.N., Inoue, R., Mori, H., Billard, J.-M., et al. (2013). Neuronal d-Serine and Glycine Release Via the Asc-1 Transporter Regulates NMDA Receptor-Dependent Synaptic Activity. J. Neurosci. 33, 3533–3544. 10.1523/JNEUROSCI.3836-12.2013.

38. Ribeiro, C.S., Reis, M., Panizzutti, R., de Miranda, J., and Wolosker, H. (2002). Glial transport of the neuromodulator d-serine. Brain Res. 929, 202–209. 10.1016/S0006-8993(01)03390-X.

39. Kory, N., Wyant, G.A., Prakash, G., uit de Bos, J., Bottanelli, F., Pacold, M.E., Chan, S.H., Lewis, C.A., Wang, T., Keys, H.R., et al. (2018). SFXN1 is a mitochondrial serine transporter required for one-carbon metabolism. Science 362, eaat9528. 10.1126/science.aat9528.

40. Chidley, C., Darnell, A.M., Gaudio, B.L., Lien, E.C., Barbeau, A.M., Heiden, M.G.V., and Sorger, P.K. (2023). A CRISPRi/a screening platform to study cellular nutrient transport in diverse microenvironments. 2023.01.26.525375. 10.1101/2023.01.26.525375.

41. Brown, T.C., Murtha, T.D., Rubinstein, J.C., Korah, R., and Carling, T. (2018). SLC12A7 alters adrenocortical carcinoma cell adhesion properties to promote an aggressive invasive behavior. Cell Commun. Signal. 16, 27. 10.1186/s12964-018-0243-0.

42. Zhou, X., Jiang, M., Liu, Z., Xu, M., Chen, N., Wu, Z., Gu, C., Chin, E., and Yang, X. (2021). Na^+^/H^+^-Exchanger Family as Novel Prognostic Biomarkers in Colorectal Cancer. J. Oncol. 2021, e3241351. 10.1155/2021/3241351.

43. Kanai, Y., and Hediger, M.A. (2004). The glutamate/neutral amino acid transporter family SLC1: molecular, physiological and pharmacological aspects. Pflüg. Arch. 447, 469–479. 10.1007/s00424-003-1146-4.

44. Xia, J., Zhang, J., Wu, X., Du, W., Zhu, Y., Liu, X., Liu, Z., Meng, B., Guo, J., Yang, Q., et al. (2022). Blocking glycine utilization inhibits multiple myeloma progression by disrupting glutathione balance. Nat. Commun. 13, 4007. 10.1038/s41467-022-31248-w.

45. van Geldermalsen, M., Wang, Q., Nagarajah, R., Marshall, A.D., Thoeng, A., Gao, D., Ritchie, W., Feng, Y., Bailey, C.G., Deng, N., et al. (2016). ASCT2/SLC1A5 controls glutamine uptake and tumour growth in triple-negative basal-like breast cancer. Oncogene 35, 3201–3208. 10.1038/onc.2015.381.

46. Morotti, M., Bridges, E., Valli, A., Choudhry, H., Sheldon, H., Wigfield, S., Gray, N., Zois, C.E., Grimm, F., Jones, D., et al. (2019). Hypoxia-induced switch in SNAT2/SLC38A2 regulation generates endocrine resistance in breast cancer. Proc. Natl. Acad. Sci. U. S. A. 116, 12452–12461. 10.1073/pnas.1818521116.

47. Sniegowski, T., Korac, K., Bhutia, Y.D., and Ganapathy, V. (2021). SLC6A14 and SLC38A5 Drive the Glutaminolysis and Serine–Glycine–One-Carbon Pathways in Cancer. Pharmaceuticals 14, 216. 10.3390/ph14030216.

48. Zhang, Z., Liu, R., Shuai, Y., Huang, Y., Jin, R., Wang, X., and Luo, J. (2020). ASCT2 (SLC1A5)-dependent glutamine uptake is involved in the progression of head and neck squamous cell carcinoma. Br. J. Cancer 122, 82–93. 10.1038/s41416-019-0637-9.

49. Wang, Q., Hardie, R.-A., Hoy, A.J., van Geldermalsen, M., Gao, D., Fazli, L., Sadowski, M.C., Balaban, S., Schreuder, M., Nagarajah, R., et al. (2015). Targeting ASCT2-mediated glutamine uptake blocks prostate cancer growth and tumour development. J. Pathol. 236, 278–289. 10.1002/path.4518.

50. Cormerais, Y., Massard, P.A., Vucetic, M., Giuliano, S., Tambutté, E., Durivault, J., Vial, V., Endou, H., Wempe, M.F., Parks, S.K., et al. (2018). The glutamine transporter ASCT2 (SLC1A5) promotes tumor growth independently of the amino acid transporter LAT1 (SLC7A5). J. Biol. Chem. 293, 2877–2887. 10.1074/jbc.RA117.001342.

51. Bröer, A., Brookes, N., Ganapathy, V., Dimmer, K.S., Wagner, C.A., Lang, F., and Bröer, S. (1999). The astroglial ASCT2 amino acid transporter as a mediator of glutamine efflux. J. Neurochem. 73, 2184–2194.

52. Fu, T.-F., Rife, J.P., and Schirch, V. (2001). The Role of Serine Hydroxymethyltransferase Isozymes in One-Carbon Metabolism in MCF-7 Cells as Determined by 13C NMR. Arch. Biochem. Biophys. 393, 42–50. 10.1006/abbi.2001.2471.

53. Tedeschi, P.M., Markert, E.K., Gounder, M., Lin, H., Dvorzhinski, D., Dolfi, S.C., Chan, L.L.-Y., Qiu, J., DiPaola, R.S., Hirshfield, K.M., et al. (2013). Contribution of serine, folate and glycine metabolism to the ATP, NADPH and purine requirements of cancer cells. Cell Death Dis. 4, e877–e877. 10.1038/cddis.2013.393.

54. Hennequart, M., Labuschagne, C.F., Tajan, M., Pilley, S.E., Cheung, E.C., Legrave, N.M., Driscoll, P.C., and Vousden, K.H. (2021). The impact of physiological metabolite levels on serine uptake, synthesis and utilization in cancer cells. Nat. Commun. 12, 6176. 10.1038/s41467-021-26395-5.

55. Oh, D.S., Troester, M.A., Usary, J., Hu, Z., He, X., Fan, C., Wu, J., Carey, L.A., and Perou, C.M. (2006). Estrogen-Regulated Genes Predict Survival in Hormone Receptor–Positive Breast Cancers. J. Clin. Oncol. 24, 1656–1664. 10.1200/JCO.2005.03.2755.

56. Budczies, J., Brockmöller, S.F., Müller, B.M., Barupal, D.K., Richter-Ehrenstein, C., Kleine-Tebbe, A., Griffin, J.L., Orešič, M., Dietel, M., Denkert, C., et al. (2013). Comparative metabolomics of estrogen receptor positive and estrogen receptor negative breast cancer: alterations in glutamine and beta-alanine metabolism. J. Proteomics 94, 279–288. 10.1016/j.jprot.2013.10.002.

57. Jia, M., Andreassen, T., Jensen, L., Bathen, T.F., Sinha, I., Gao, H., Zhao, C., Haldosen, L.-A., Cao, Y., Girnita, L., et al. (2016). Estrogen Receptor α Promotes Breast Cancer by Reprogramming Choline Metabolism. Cancer Res. 76, 5634–5646. 10.1158/0008-5472.CAN-15-2910.

58. Wang, C.-Y., Chiao, C.-C., Phan, N.N., Li, C.-Y., Sun, Z.-D., Jiang, J.-Z., Hung, J.-H., Chen, Y.-L., Yen, M.-C., Weng, T.-Y., et al. (2020). Gene signatures and potential therapeutic targets of amino acid metabolism in estrogen receptor-positive breast cancer. Am. J. Cancer Res. 10, 95–113.

59. Cantor, J.R., Abu-Remaileh, M., Kanarek, N., Freinkman, E., Gao, X., Louissaint, A., Lewis, C.A., and Sabatini, D.M. (2017). Physiologic Medium Rewires Cellular Metabolism and Reveals Uric Acid as an Endogenous Inhibitor of UMP Synthase. Cell 169, 258–272.e17. 10.1016/j.cell.2017.03.023.

60. Missiaen, R., Anderson, N.M., Kim, L.C., Nance, B., Burrows, M., Skuli, N., Carens, M., Riscal, R., Steensels, A., Li, F., et al. (2022). GCN2 inhibition sensitizes arginine-deprived hepatocellular carcinoma cells to senolytic treatment. Cell Metab. 34, 1151–1167.e7. 10.1016/j.cmet.2022.06.010.

61. Locasale, J.W. (2013). Serine, glycine and one-carbon units: cancer metabolism in full circle. Nat. Rev. Cancer 13, 572–583. 10.1038/nrc3557.

62. Maddocks, O.D.K., Athineos, D., Cheung, E.C., Lee, P., Zhang, T., van den Broek, N.J.F., Mackay, G.M., Labuschagne, C.F., Gay, D., Kruiswijk, F., et al. (2017). Modulating the therapeutic response of tumours to dietary serine and glycine starvation. Nature 544, 372–376. 10.1038/nature22056.

63. Wang, W., Pan, H., Ren, F., Chen, H., and Ren, P. (2022). Targeting ASCT2-mediated glutamine metabolism inhibits proliferation and promotes apoptosis of pancreatic cancer cells. Biosci. Rep. 42, BSR20212171. 10.1042/BSR20212171.

64. van Geldermalsen, M., Quek, L.-E., Turner, N., Freidman, N., Pang, A., Guan, Y.F., Krycer, J.R., Ryan, R., Wang, Q., and Holst, J. (2018). Benzylserine inhibits breast cancer cell growth by disrupting intracellular amino acid homeostasis and triggering amino acid response pathways. BMC Cancer 18, 689. 10.1186/s12885-018-4599-8.

65. Chiu, M., Sabino, C., Taurino, G., Bianchi, M.G., Andreoli, R., Giuliani, N., and Bussolati, O. (2017). GPNA inhibits the sodium-independent transport system l for neutral amino acids. Amino Acids 49, 1365–1372. 10.1007/s00726-017-2436-z.

66. Schulte, M.L., Fu, A., Zhao, P., Li, J., Geng, L., Smith, S.T., Kondo, J., Coffey, R.J., Johnson, M.O., Rathmell, J.C., et al. (2018). Pharmacological blockade of ASCT2-dependent glutamine transport leads to antitumor efficacy in preclinical models. Nat. Med. 24, 194–202. 10.1038/nm.4464.

67. Bröer, A., Fairweather, S., and Bröer, S. (2018). Disruption of Amino Acid Homeostasis by Novel ASCT2 Inhibitors Involves Multiple Targets. Front. Pharmacol. 9.

68. Garaeva, A.A., Oostergetel, G.T., Gati, C., Guskov, A., Paulino, C., and Slotboom, D.J. (2018). Cryo-EM structure of the human neutral amino acid transporter ASCT2. Nat. Struct. Mol. Biol. 25, 515–521. 10.1038/s41594-018-0076-y.

69. Yu, X., Plotnikova, O., Bonin, P.D., Subashi, T.A., McLellan, T.J., Dumlao, D., Che, Y., Dong, Y.Y., Carpenter, E.P., West, G.M., et al. (2019). Cryo-EM structures of the human glutamine transporter SLC1A5 (ASCT2) in the outward-facing conformation. eLife 8, e48120. 10.7554/eLife.48120.

70. Kruse, T., Reiber, H., and Neuhoff, V. (1985). Amino acid transport across the human blood-CSF barrier: An evaluation graph for amino acid concentrations in cerebrospinal fluid. J. Neurol. Sci. 70, 129–138. 10.1016/0022-510X(85)90082-6.

71. Dolgodilina, E., Imobersteg, S., Laczko, E., Welt, T., Verrey, F., and Makrides, V. (2016). Brain interstitial fluid glutamine homeostasis is controlled by blood-brain barrier SLC7A5/LAT1 amino acid transporter. J. Cereb. Blood Flow Metab. Off. J. Int. Soc. Cereb. Blood Flow Metab. 36, 1929–1941. 10.1177/0271678X15609331.

72. Rainesalo, S., Keränen, T., Palmio, J., Peltola, J., Oja, S.S., and Saransaari, P. (2004). Plasma and cerebrospinal fluid amino acids in epileptic patients. Neurochem. Res. 29, 319–324. 10.1023/b:nere.0000010461.34920.0c.

73. Maggs, D.G., Jacob, R., Rife, F., Lange, R., Leone, P., During, M.J., Tamborlane, W.V., and Sherwin, R.S. (1995). Interstitial fluid concentrations of glycerol, glucose, and amino acids in human quadricep muscle and adipose tissue. Evidence for significant lipolysis in skeletal muscle. J. Clin. Invest. 96, 370–377. 10.1172/JCI118043.

74. Ngo, B., Kim, E., Osorio-Vasquez, V., Doll, S., Bustraan, S., Liang, R.J., Luengo, A., Davidson, S.M., Ali, A., Ferraro, G.B., et al. (2020). Limited Environmental Serine and Glycine Confer Brain Metastasis Sensitivity to PHGDH Inhibition. Cancer Discov. 10, 1352–1373. 10.1158/2159-8290.CD-19-1228.

75. Palmer, A.C., Chidley, C., and Sorger, P.K. (2019). A curative combination cancer therapy achieves high fractional cell killing through low cross-resistance and drug additivity. eLife 8, e50036. 10.7554/eLife.50036.

76. Stender, J.D., Nwachukwu, J.C., Kastrati, I., Kim, Y., Strid, T., Yakir, M., Srinivasan, S., Nowak, J., Izard, T., Rangarajan, E.S., et al. (2017). Structural and Molecular Mechanisms of Cytokine-Mediated Endocrine Resistance in Human Breast Cancer Cells. Mol. Cell 65, 1122–1135.e5. 10.1016/j.molcel.2017.02.008.

77. Kang, Y.P., Torrente, L., Falzone, A., Elkins, C.M., Liu, M., Asara, J.M., Dibble, C.C., and DeNicola, G.M. (2019). Cysteine dioxygenase 1 is a metabolic liability for non-small cell lung cancer. eLife 8, e45572. 10.7554/eLife.45572.

78. Kang, Y.P., Mockabee-Macias, A., Jiang, C., Falzone, A., Prieto-Farigua, N., Stone, E., Harris, I.S., and DeNicola, G.M. (2021). Non-canonical Glutamate-Cysteine Ligase Activity Protects against Ferroptosis. Cell Metab. 33, 174–189.e7. 10.1016/j.cmet.2020.12.007.

